# Fast and Accurate Kinship Estimation Using Sparse SNPs in Relatively Large Database Searches

**DOI:** 10.1101/2022.08.22.504804

**Authors:** June Snedecor, Tim Fennell, Seth Stadick, Nils Homer, Joana Antunes, Kathryn Stephens, Cydne Holt

## Abstract

Forensic genetic genealogy (FGG) has primarily relied upon dense single nucleotide polymorphism (SNP) profiles from forensic samples or unidentified human remains queried against online genealogy database(s) of known profiles generated with SNP microarrays or from whole genome sequencing (WGS). In these queries, SNPs are compared to database samples by locating contiguous stretches of shared SNP alleles that allow for detection of genomic segments that are identical by descent (IBD) among biological relatives (kinship). This segment-based approach, while robust for detecting distant relationships, generally requires DNA quantity and/or quality that are sometimes not available in forensic casework samples. By focusing on SNPs with maximal discriminatory power and using an algorithm designed for a sparser SNP set than those from microarray typing, performance similar to segment matching was reached even in difficult casework samples. This algorithm locates shared segments using kinship coefficients in “windows” across the genome. The windowed kinship algorithm is a modification of the PC-AiR and PC-Relate tools for genetic relatedness inference, referred to here as the “whole genome kinship” approach, that control for the presence of unknown or unspecified population substructure. Simulated and empirical data in this study, using DNA profiles comprised of 10,230 SNPs (10K multiplex) targeted by the ForenSeq™ Kintelligence Kit demonstrate that the windowed kinship approach performs comparably to segment matching for identifying first, second and third degree relationships, reasonably well for fourth degree relationships, and with fewer false kinship associations. Selection criteria for the 10K SNP PCR-based multiplex and functionality of the windowed kinship algorithm are described.

## 2. Introduction

Forensic genetic genealogy (FGG), also known as investigative genetic genealogy (IGG), refers to investigative lead generation using dense single nucleotide polymorphism (SNP) profiles from unidentified human remains or crime scene samples that are queried against direct-to-consumer (DTC) genealogical database(s) comprised of known, reference SNP profiles to associate with various degree relatives. FGG has gained interest from the forensic and law enforcement community as a tool to consider when CODIS searching and other means have been exhausted [1]. A large SNP profile generated from microarray analysis is expected to have better discriminatory power than the current battery of forensically relevant short tandem repeat (STR) loci. However, microarray technology requires DNA of quality and input [2] that may not be available from crime scenes or human remains such as skeletal remains.

In addition, for investigative purposes, identifying more distant relationship matches can require a substantially higher effort than for closer relationships. Fig. 1 shows example relationships out to seventh degree. With each increase in degree the number of possible family trees increases significantly. Many genealogy investigations focus on third degree or closer relationships due to burden and inefficiencies that can occur when distance extends to fourth degree or beyond [3]. A polymerase chain reaction (PCR) based FGG typing system that targets sufficient kinship SNPs with high sensitivity of detection of first, second and third degrees relatives, and good detection of fourth and fifth degree relationships, can assist to address the technology gap between microarray and WGS SNP methods regarding sample quality and quantity, personal health information, time and cost.

**Fig. 1:**
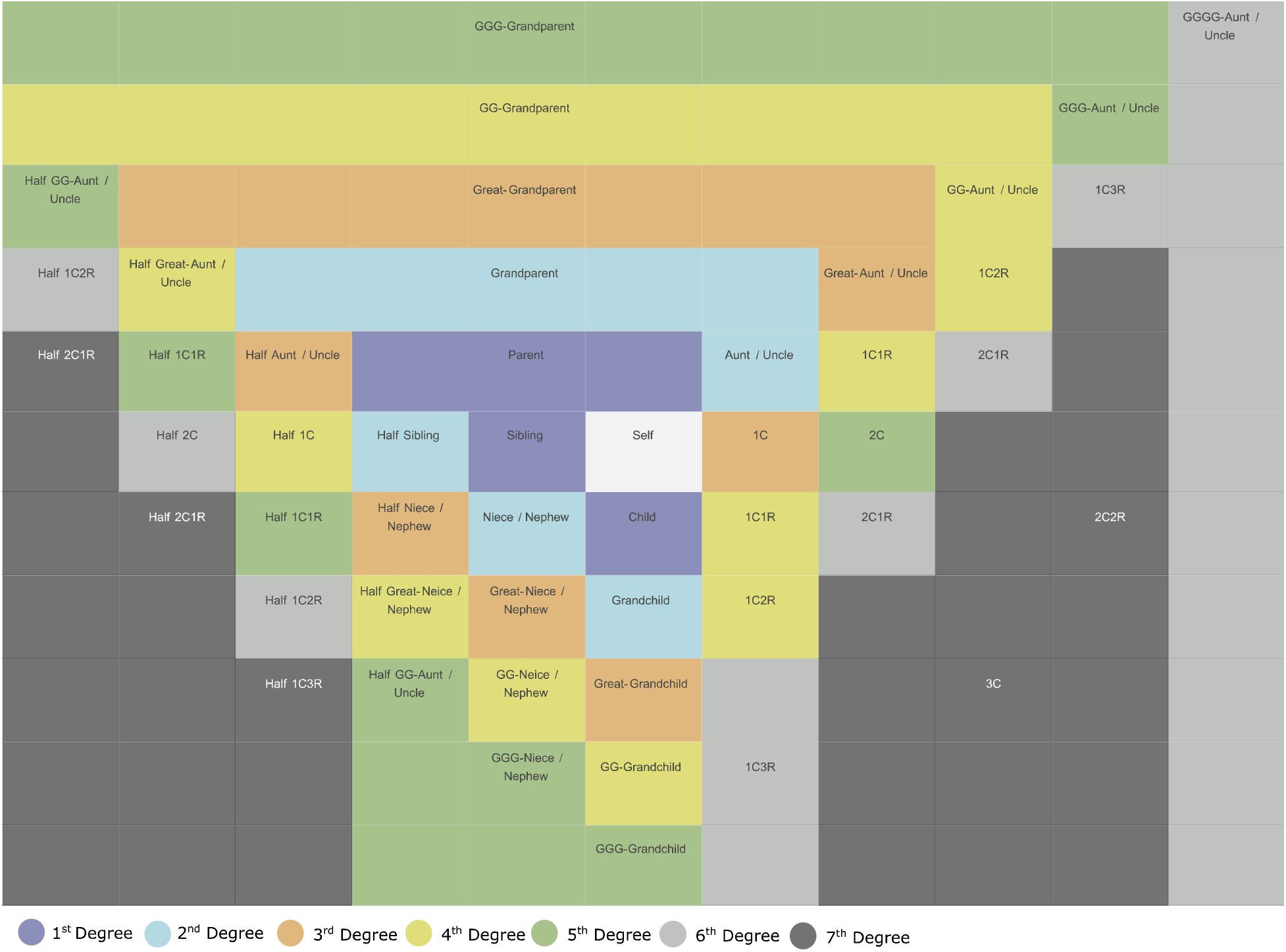
Examples of degrees of human genetic relationships (adapted from DNA Painter, https://dnapainter.com/).

Generally, FGG has used a “segment matching” approach to estimate kinship by finding contiguous blocks (usually numbering in the hundreds) of identical shared alleles and estimating the total centimorgan (cM) distance covered by those segments [4][5][6]. Segment matching across the genome requires many hundreds of thousands of SNPs thus the use of microarrays or WGS on forensic samples. SNPs that are physically linked on a chromosome are more likely to be inherited together (identical by descent (IBD)), therefore much of the information used in segment matching is redundant and intentionally so. However, as is the case with identity by state (IBS) methods, fewer SNPs can be successfully used for sensitive and specific kinship detection when they provide enough information [7]. A SNP hybridization capture panel used a similar approach using a limited SNP panel [8] and also explored alternative approaches to segment matching in order to evaluate kinship [9].

Forensic genetics has relied upon PCR for decades and can be used to target kinship informative SNPs for FGG. A targeted, forensic PCR assay and analytical software that recovers SNP allele calls from low level, damaged and/or partially degraded forensic DNA samples in a manner sufficient for FGG query was developed. With this strategy, DNA sample analyses may be conducted in operational laboratories using desktop sequencers followed by genealogical database query using a companion kinship inference method. This study describes selection criteria for 10,230 high value SNPs targeted by the ForenSeq Kintelligence™ Kit (Verogen, Inc., San Diego CA), referred to here as the 10K multiplex, and a windowed kinship algorithm to accurately locate and classify kinship out to fourth degree relatives. Of these loci, 9,867 are kinship informative SNPs selected from the Infinium CytoSNP-850K BeadChip and Global Screening Array (Illumina, Inc., San Diego, CA) and filtered using the Genome Aggregation Database (gnomAD) v3.0, the Single Nucleotide Polymorphism database (dbSNP) v151 and GEDmatch for robust representation across global populations. The SNPs are maximally spaced across the genome to minimize linkage effects and have no reported significance in ClinVar [10] (Fig. 2). The remaining 363 SNPs can be used to inform biogeographical ancestry, identity, hair and eye color, or biological sex. Identity SNPs were included in order to allow cross checking of kinship using a previously validated assay (ForenSeq DNA Signature™). The companion windowed kinship algorithm was built upon PC-AiR [11][7][12] and PC-Relate [13] methods, referred to here as the whole genome kinship method, with an additional windowing component. This windowed kinship algorithm also relies on the concept of segment matching (*i.e*., that distant relatives share contiguous blocks of identical SNPs) and locates segments as blocks of highly scored kinship rather than stretches of identical SNP allele calls to provide even higher performance for FGG.

**Fig. 2:**
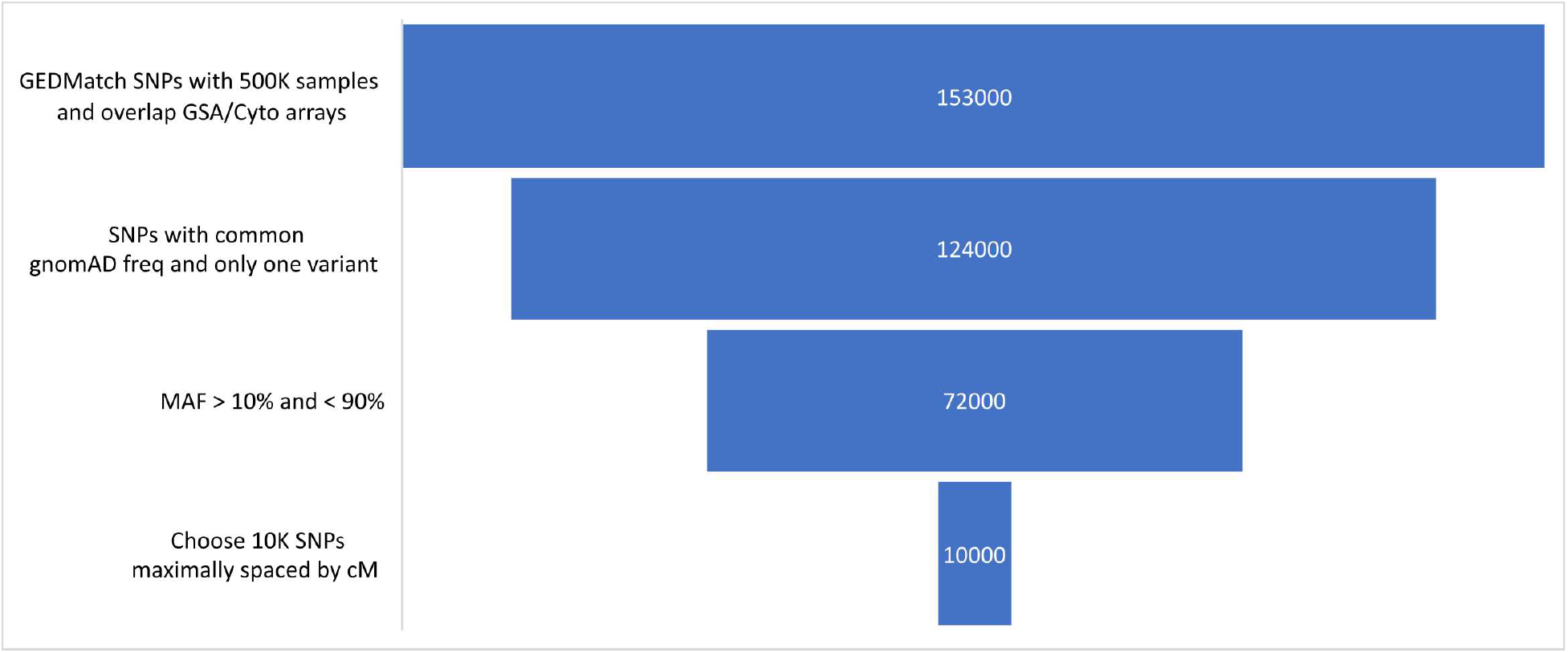
Method for kinship SNP selection. The overall selection strategy as well as the number of SNPs that remained after each stage of filtering are shown. SNPs that were well represented among DTC microarrays were prioritized then limited to SNPs with gnomAD European allele frequencies that were approximated (within three fold) those observed in the GEDmatch database. SNPs with minor allele frequencies (MAF) < 10% or > 90% were excluded. The resulting 72,000 SNPs were evaluated using the windowed kinship algorithm. 9,867 maximally spaced (cM) kinship informative SNPs were optimized in a PCR-based multiplex.

Simulated pedigrees and real microarray profiles from GEDmatch were used to assess performance of the windowed algorithm. Additionally, two known pedigrees were analyzed using the ForenSeq Kintelligence Kit to assess further kinship estimation performed on real DNA samples using the windowed kinship algorithm. To use GEDmatch microarray profiles as knowns for true relationships, expected degrees of relationship were set using segment matching information since multiple, real extended pedigrees were not available. The 10K SNPs for the 10K FGG multiplex were selected from the GEDmatch test set and the windowed kinship approach was compared to the PC-AiR/PC-Relate whole genome kinship method out to fifth degree relationships.

## 3. Materials and Methods

### 3.1. SNP Reference Data for Algorithm Testing

1000 anonymized query samples were selected at random from GEDmatch^1^, to generate a test set of SNP profiles with varying degrees of relationship (see Fig. S1 for country of origin for test set and Fig. S2 for GEDmatch country of origin). For each query sample a single sample (if found) was selected for a set of varying total shared cM ranges calculated by the GEDmatch one-to-many tool (2787-3600, 1083-2787, 326-1083 and 0-326 cM) and was added to the target set. A target set of 2,954 samples (including the original set of 1000 query samples) was compiled. Since most donors of GEDmatch samples are unrelated, the test set was developed to ensure that there was a sufficient set of related samples representing each degree of interest (Table S1).

To evaluate sensitivity and specificity within a particular degree of relationship, the test set was filtered to all sample pairs with the expected shared cM range for that relationship (as shown in Table S1) and all pairs that had zero shared cM. Pairs of samples that had fewer than 9,000 mutually called loci in the 10K SNP multiplex (see Section 4.1 for details) were not considered (see Fig. S3 for overlapping loci counts in the test set). See Table S2 for sample pair totals for different relationship levels.

For simulated pedigrees, genotype data from the 1000 Genomes Project (1KGP) (Phase 3 build 20130502) [14][15] were used as pedigree founders. The original set of 2,504 samples was then filtered to remove relatives using the windowed kinship method and the 10K SNP loci. Sample pairs with > 100 shared total cM and a longest shared segment >30 cM were removed, which reduced the set to 1,851 founder samples. Ped-sim [16] was used to generate 200 pedigrees from these founders using the Poisson model and sex averaged map from Bhérer *et al*. [17]. Relationships were simulated as follows: sibling (first degree), half sibling (second degree), first cousin (third degree), half cousin (fourth degree) and second cousin (fifth degree) (see Fig. S7 for pedigrees). For each relationship degree, there were 200 true relationships and 79,600 unrelated pairings from 400 total samples. To determine how many matching SNPs could be expected for each relationship degree, 1000 independent sample pairs per degree were generated, some with overlapping founder samples. Pairs that shared founders were not compared to each other.

Two known, extended pedigrees were also used to test the 10K SNPs and the windowed kinship algorithm. All samples obtained for testing with Kintelligence were obtained after volunteers signed an informed consent form authorizing the use of de-identified samples for research use publication. One pedigree (n = 26 individuals) included relatives out to the sixth degree (see Fig. S8) uploaded on the public GEDmatch database. Relatives in GEDmatch were marked with their known relationships and anonymized. Since profiles on genealogy databases have been generated by different arrays over time, this evaluation provided a real-world example of performance on DTC data. A “self” reference buccal sample (V024) was typed using the ForenSeq Kintelligence kit, MiSeq FGx sequencer and Universal Analysis Software 2.6, and kinship analysis was performed against the entire GEDmatch database. The second pedigree (n = 15 individuals) was generated from gDNA from buccal swabs for relatives out to the fifth degree (see Fig. S10) typed with the ForenSeq Kintelligence kit.

The V004, V016, V017, V018, V019, V020, V021, and V024 samples consisted of contemporary buccal swabs extracted with the QIAamp DNA Investigator kit (Qiagen, CA), according to the manufacturer’s instructions. DNA quantification was performed using the Quantifluor^®^ ONE dsDNA System (Promega, WI). To degrade V016, 16.8 ng of DNA was placed in each of 4 PCR tubes. All 4 replicates were subjected to continuous cycles of 98 °C for 1 hour, and 4 °C for 10 min for 24 hours, followed by an indefinite 4 °C hold. The DNA replicates were then centrifuged in a tabletop centrifuge for 1 min at maximum speed in order to concentrate any liquid particles to the bottom of the tube. To maximize recovery of DNA, 15 μL of water was used for each replicate, by pipetting the sides of the tubes 10 times, followed by vortexing and centrifugation to allow the DNA samples to be collected in the bottom of the tubes. Samples were quantified using the Quantifluor^®^ ONE dsDNA (Promega, WI). To simulate ante-mortem samples, 1 ng input of each sample was amplified using the ForenSeq™ Kintelligence kit. To simulate post-mortem samples, degraded and/or low input (< 1 ng) samples were amplified using the ForenSeq™ Kintelligence kit, according to the manufacturer’s instructions, in following manner: the degraded V016 replicate was amplified with 1 ng input, and the V016 replicate amplified with 250 pg input. All libraries were sequenced using the MiSeq FGx™ reagent kit and the MiSeq FGx™ instrument.

### 3.2. 10K SNP PCR-Based Multiplex Design

To maximize the value of SNPs in the Kintelligence multiplex, locus selection criteria were considered (see Fig. 2). First, SNPs were selected that are well represented in genetic genealogy databases like GEDmatch. As a quality control measure the frequencies represented in GEDmatch were assessed for general agreement (within three fold of) the population frequencies reported by the Genome Aggregation Database (gnomAD) for European ancestry [18] since this group represents the geographical location of majority of samples in GEDmatch. SNPs were selected that have demonstrated variability within all major human subpopulations (see S2_gnomad.genomes.r3.0.kintelligence_filtered.vcf.zip for population frequencies). With a rare allele at a biallelic SNP, most individuals will be homozygous for the reference allele which is not generally informative for kinship inference in large databases. Common SNP alleles increase the chances for informative differences and similarities between samples. SNPs designated as benign/likely benign in ClinVar were selected [10]. No SNPs with any clinical significance in ClinVar were included. 9,867 kinship informative SNPs that met the selection criteria were included in the ForenSeq Kintelligence multiplex design and are maximally spaced (cM) along each autosome to minimize the effects of physical linkage thereby maximizing the informational value of each individual locus (see Fig. S5 for cM distances in final multiplex).

The kinship informative SNP selection method (Fig. 2) was as follows:

1. Find intersection of SNP content on Infinium Global Screening Array (GSA) and Infinium CytoSNP-850K (Cyto).
2. Filter SNPs in GEDmatch to those in the GSA and Cyto list.
3. Keep SNPs with > 500,000 profiles in GEDmatch (^~^40% of the GEDmatch database at the time the 10K multiplex was designed in July 2020)
4. Filter to SNPs in gnomAD where GEDmatch frequency is within three-fold of gnomAD EUR allele frequency. This is intended to only capture gross discrepancies between gnomAD and GEDmatch, the frequencies can be quite divergent and still be considered (i.e., 16% gnomAD and 45% GEDmatch would still be included, while 14% to 45% would not.)
  - Enforce gnomAD minor allele frequency (MAF) frequency between 0-50% since some “minor alleles” in gnomAD are actually the major allele. Functionally, this means that if a SNPs reported frequency is > 50%, we use the frequency 100-reported frequency
  - Calculate ratio between GEDmatch and gnomAD if GEDmatch is larger or vice versa
  - If ratio >= 3, discard
5. Remove loci with more than one gnomAD SNP within three-fold of the GEDmatch frequency. This is both because GEDmatch only retains one allele per locus, and because genotypes from arrays may be untrustworthy in triallelic situations. For example, a microarray which is probing for A and C may call an A/G as A/A.
6. Remove SNPs where a population (GEDmatch or nine subpopulations in gnomAD in Table S3) have MAF < 10% or > 90%.
7. Choose N SNPs from the remaining set as follows:
  - Divide N among autosomes relative to their length in cM
  - Compute average spacing for each chromosome in cM
  - Window across the chromosome as follows:
    1. Find next SNP on the chromosome
    2. Pull all SNPs within 70% of the average cM spacing
    3. Pick SNP with the most samples in GEDmatch and the MAF closest to 50%
    4. Discard SNPs within 30% of the average cM spacing downstream of the chosen SNP
    5. Repeat

### 3.3. Statistical Methods

#### 3.3.1. PC-AiR with Modified Unrelated Set Selection

Model based ancestry estimation methods are less accurate in the presence of genetic relatedness as they cannot distinguish between ancestral groups and clusters of more recent relatives [19]. The PC-AiR [11] method consists of two steps: 1) select a maximally ancestrally diverse set of unrelated samples from a source set; and 2) perform principal component analysis (PCA) on the ancestry representative subset and predict components of variation for all remaining individuals based on genetic similarities. PC-AiR defines a method for identifying a set of unrelated samples that works well for modest sample sets but does not scale well. In a database with *n* samples, the algorithm must perform *n^2^* comparisons to remove each related sample. For smaller datasets, this approach is acceptable to maximize ancestral divergence. For relatively large databases a pairwise comparison approach becomes infeasible. Consider, if a database of 1.5M has a thousand related samples then (1.5 million)^2^ * 1000 or 2.25 * 10^15^ calculations are required.

Alternatively, relatives can be assessed, and samples discarded to generate an “unrelated” sample set, which can be searched in a much less computationally demanding fashion. Beginning with samples that have the fewest total related samples to minimize data loss, samples can be added iteratively to the unrelated set while relatives are immediately removed from consideration. A more stringent kinship statistic can also be used to find relatives under the assumption that since there is a larger initial dataset, removal of more potential relatives from consideration can be tolerated and helps to ensure that the final set does not contain relatives or if so minimally. Also, samples with a high number (>9000 for the 10K multiplex) of called loci can be considered in the chosen multiplex. The following algorithm was developed as a modification of the PC-AiR method:

1. Remove all samples with >= 5% missing data from the SNP set being used.
2. Compute KING-Robust [11] kinship coefficient between all pairs of samples *N*. This kinship coefficient for individuals *i* and *j* are denoted as *φ_ij_* and is defined as the probability that a random allele selected from *i* and a random allele selected from *j* at a locus are identical by descent (IBD). Use a relatedness threshold *τ*_*φ*1_= 0.01 to determine whether the pair of samples are expected to be IBD. Use a relatedness threshold of *τ*_*φ*2_= 0.025 for ancestry divergent samples.
  a. Call *φ_ij_* with kinship coefficient > *τ*_*φ*1_ as related
  b. Call *φ_ij_* with kinship coefficient < −*τ*_*φ*2_ as ancestrally divergent.
3. Initialize two subsets *U* = *∅* and *R* = ∅ where ∅ is the empty set.
4. For all *i*∈*N*
  a. *r_i_* = the set of all *j* where *φ_ij_* > *τ*_*φ*1_ for *j* ∈ *N* and *j* ≠ *i*. *r_i_* is the set of all relatives for *i*.
  b. *d_i_* = the set of all *j* where *φ_ij_* < – *τ*_*φ*2_ for *j* ∈ *N* and *j* ≠ *i*. *d_i_* is the set of ancestrally divergent relatives for *i*
5. Rank all samples *i∈N* by |*r_i_*| in ascending order.
6. For samples in *N* with the same |*r*|, sort by |*d*| in descending order.
7. Iterate through ranked samples and for *i ∈ N*
  a. If *i* ∉ *R*, *U* = *U* ∪ *i* and *R* = *R* U ∪ *r_i_*.
  b. If *i* ∈ *R* continue to next iteration.

Calculating pairwise kinships is still O(π^2^), however the windowed kinship algorithm performs that step only once per model build instead of after every removal of a relative as in the unmodified PC-AiR. Once the unrelated set has been determined, the principal components are determined from the set *U* using the original PC-AiR method.

#### 3.3.2. PC-Relate and Windowed PC-Relate

Many current methods for kinship inference either assume that pairs of samples came from a homogenous population or require that samples be categorized by sub-population. PC-Relate [13] uses principal components from PC-AiR and partitions genetic correlations into two separate components: a component for the sharing of alleles that are IBD from recent common ancestors and another component for allele sharing due to more distant common ancestry.

Assuming the top PC components from PC-AiR correctly capture the population structure of the samples, those components can be used to estimate the expected allele frequencies based on an individual’s ancestral background using a linear regression model rather than using a static population frequency. As described by Conomos *et al*. regarding PC-Relate [13] for a particular SNP s and an individual *i*, 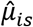 can be calculated which represents the specific expected population SNP frequency for this individual’s background as a substitute for 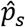 which is simply the global expected frequency for that SNP determined from a population database.

Once the SNP frequencies have been estimated for each individual it is straightforward to estimate the kinship coefficient *ϕ_ij_* for individuals *i* and *j* for a set of SNPs *S*. Let *g_is_* be the number of reference alleles an individual has at SNP s.

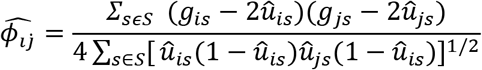

The estimator 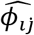 measures the scaled residual genetic covariance between *i* and *j* after conditioning on their respective ancestries. Overall, this measurement of kinship can work well. For the FGG use case it has limitations at distant relationships. With the 10K SNP multiplex, the expected number of IBD SNPs for a fifth degree relationship is approximately 300, assuming approximately 0.3 cM between SNPs and 100 total shared cM. Thus, even random fluctuations in overall allele sharing can be above the threshold for detecting a distant relative, which was clear when comparing GEDmatch segment matching against the whole genome kinship coefficient at more distant degrees of relationship.

It is well understood that physically linked genomic regions are more likely to be from inherited DNA which is clustered in contiguous blocks that are reduced in size with each generation. Conversely random allele sharing is in general spread throughout the genome. Segment matching used in GEDmatch and Ancestry.com [20], rely upon this basic concept. A similar approach was taken here by calculating “windows” of kinship across the genome to find shared kinship segments and boost specificity in estimating the more distant relationships.

Given a set of SNPs *S* = {*s*_0_..*s_n_*} and a window size *l* a window of SNPs is defined at index *k* as *w_k_* = {*s_k_*.. *s*_*k*+l_}. The windowed kinship approach is as follows:

1. Enumerate all possible windows *W* = {*w*_0_.. *w*_|*S*|–*l*_}· Windows must be contained within a single chromosome.
2. Given an individual *i* and an individual *j*
3. Calculate kinship across all windows. For *k* = {0.. |*W*|}

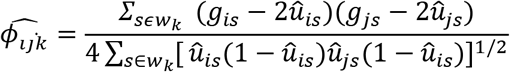

From here locate IBD segments as follows:

1. Create an empty set *P* to contain all windows with kinship above threshold *t*.
2. Given an individual *i* and an individual *j*
  a. Iterate through *k* = {0.. |*W*|}.
  b. If 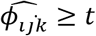 add *w_k_* to *P*
  c. If 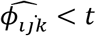 continue
3. Merge windows that have overlapping genomic positions
  a. Iterate through *P*.
  b. If the current window overlaps with the next window, remove next window from *P* and reset last index in current window with values from next.
  c. Repeat until there are no overlapping windows remaining.
4. Remove all windows where the fraction of SNPs at which the individuals *i* and *j* share at least one allele is lower than a threshold value *f* (*e.g. f=0.95*). In windows of true IBD the fraction should be 1 in the absence of genotyping error. However false kinship signal can be generated when many SNPs share no alleles but many others share both alleles.

To find total shared cM, a two-pass approach was taken, first identifying segments with stretches of SNPs with at least one shared allele (half match) and the second, within those segments, stretches of SNPs that have two shared alleles (full match). Half matching segments have *t* = 0.22 while full matching segments have *t* = 0.44. These are reduced from the theoretical values of 0.25 and 0.5 under a strict kinship definition; in the windowed kinship algorithm the thresholds are set slightly lower to allow for genotyping error. When calculating total shared cM, first degree relationships can be distinguished as they mostly consist of half matches and consanguineous or self matches and a higher degree of full matches than more distant relationships.

## 4. Results and Discussion

### 4.1. Evaluating Kinship Informative SNP Multiplex Size

Most genetic genealogy databases use a segment matching approach. Segment matching identifies long stretches of matching SNPs, relying on the fact SNPs that are IBD are inherited in contiguous physical blocks. Since large numbers of SNPs are queried, missing or incorrect SNP calls can have minimal effect on segment matching. For FGG, a 10K PCR-based SNP multiplex was designed to provide maximum kinship information with minimal locus content and without clinically relevant loci or disease markers (Fig. 2). These sparser data, as compared to microarray content, can be generated in one MiSeq FGx run but are less informative for kinship if standard segment matching were used. A companion, windowed kinship algorithm was developed that maximizes kinship resolution from the 10K SNP multiplex. This method starts with the same core concept as segment matching, namely identifying contiguous blocks of shared DNA. Then, rather than simply counting matching SNP allele calls, the kinship coefficient described in Conomos *et al*. with PC-Relate [13] is used as a criterion of genetic relatedness. By calculating kinship coefficients in windows across the genome, the discriminatory power of fewer SNPs was enhanced by controlling for background frequencies and population substructure (see Section 3.2).

As shown in Fig. 2, 72,000 SNPs met the locus selection criteria for a PCR-based FGG multiplex. Testing of multiplexes with varied SNP numbers was performed in combination with the windowed kinship algorithm in order to balance the number of SNPs with the ability to detect third degree relatives with high sensitivity. For example, a 20K SNP multiplex and the 10K multiplex were tested and compared using genotype data simulated by ped-sim on 1KGP founder samples for detection of kinship of degrees one through five. Based on the observed fractions of shared alleles from these simulated data (Fig. S4), the 10K and 20K SNP sets enabled significant separation between sample pairs representing third degree. The 10K and 20K SNP sets were then tested using the same simulated data with the windowed kinship approach directly. Out to the third degree, receiver operating characteristic (ROC) curves were nearly identical for the 10K and the 20K SNP sets (and could reach 100% for both sensitivity and specificity). Sensitivity in this instance means the percentage of total related pairs above the scoring thresholds and specificity means the percentage of unrelated samples above the scoring thresholds based on total shared cM and longest shared cM segment. A receiver operating characteristic (ROC) curve with an L shape that aligns closely to the upper left-hand corner indicates that adjusting the thresholds follows a predictable pattern and that there exists at least one threshold with 100% sensitivity and specificity (or close to it). As shown in Fig. 3, the ROC curve for the 10K SNP set achieves 98% sensitivity and 100% specificity for the fourth degree simulated data and performs less well on fifth degree simulated data. Performance with the 20K SNP set was better for fifth degree as expected but even in that case perfect performance was not achieved even in the best-case scenario of a full profile (all loci called). Lowering the threshold of the kinship coefficient will increase the sensitivity of the 10K multiplex comparable to the 20K multiplex with the expected decrease in specificity. Since this multiplex is intended for use with low input, low quality and/or degraded samples, the number of loci is a tradeoff between overall coverage and number of possible SNP calls. Clean, high-input samples already can use existing microarray technologies to provide more SNPs. The 10K SNP multiplex can be considered to provide a practical tool for generating investigative genetic leads extending into the fourth degree (*e.g*., first cousin once removed (1C1R)). After targeting the 10K SNP set using multiplex PCR, MiSeq FGx v3 sequencing reagents can produce 50M paired end reads, supporting a run configuration comprised of a negative control, a positive DNA control, and one forensic sample with up to 25M reads.

**Fig. 3:**
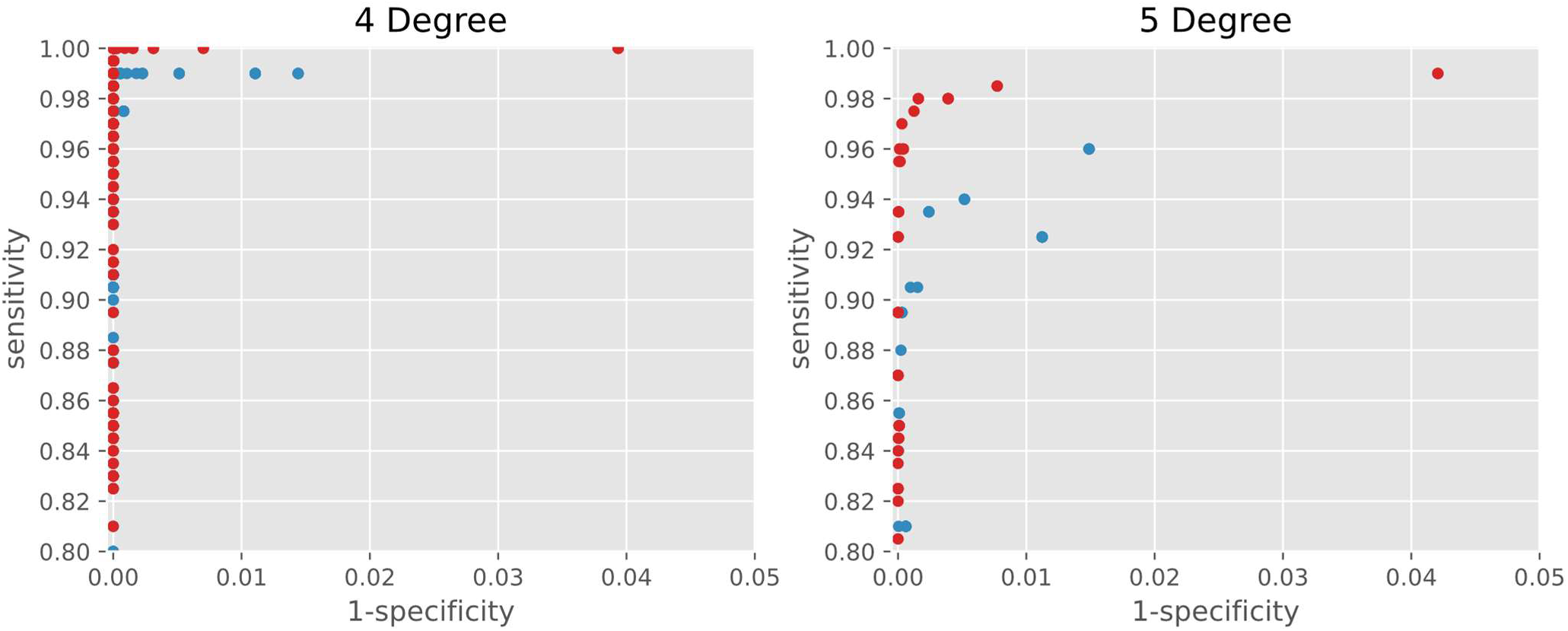
Comparison between 10K (blue dots) and 20K (red dots) SNP multiplexes of sensitivity of detection and specificity of relationship degree estimation using simulated data from ped-sim. 400 true pairs and 76,000 unrelated pairs generated per degree. For degrees one, two and three, functionally identical sensitivity and specificity were observed (100% for both sensitivity and specificity) for 10K and 20K SNPs. At fourth and fifth degree, an increase in sensitivity was observed with the 20K SNP set. Sensitivity was observed at 92.5% for 20K and 76% for 10K SNPs for fifth degree kinship, and with no false associations.

Since FGG uses relatively large databases (*i.e*., >1 million samples), evaluating the potential for false associations in the context of a list of potential candidate kinship associations can be helpful to operational settings. The term “false associations” is used here to describe pairs of samples that are above the chosen thresholds that do not actually share a relationship. These have the potential to increase operational time. As the size of a genealogy database increases, the potential also increases for unrelated sample pairs to have larger total shared cM or kinship coefficients than true relatives in a query return. As of June 2, 2022, GEDmatch contained approximately 1.5M autosomal microarray profiles^2^ and the FamilyTreeDNA database approximately 1.2M^3^. Thus, even with a specificity of 99.99%, there is potential for hundreds of candidate hits to be returned that are not actual relatives. Microarray-based DNA profiles that comprise a known pedigree extending to sixth degree relationships were used to assess limitations of the windowed kinship approach on a 1.5M sample database. One sample (V024) from the known pedigree was selected as the person of interest (“self”) and typed with the 10K multiplex (ForenSeq Kintelligence kit). The full database of 1.5 million profiles was searched using the windowed kinship algorithm described in Section 3.3 and the default thresholds implemented in the GEDmatch Pro™ (see Table S4). (Note: The GEDmatch Pro portal is dedicated to support FGG comparisons for investigative lead generations in criminal casework.) This GEDmatch test query simulated a workflow for unidentified human remains cases. All relationships out to fifth degree (2C) were detected; both sixth degree relationship pairs (2C1R) fell below the default thresholds for total shared cM, longest segment (cM) used by GEDmatch Pro for overlapping SNPs >9,000 (Table 1). Whole genome kinship is included for comparison purposes. Fifth degree kinship was associated to a synthetic profile generated from Native American genomic segments that had been uploaded to GEDmatch (confirmed by the user who initially uploaded the profile). Thus a false positive rate of 1/1,500,000 was achieved.

**Table 1:**
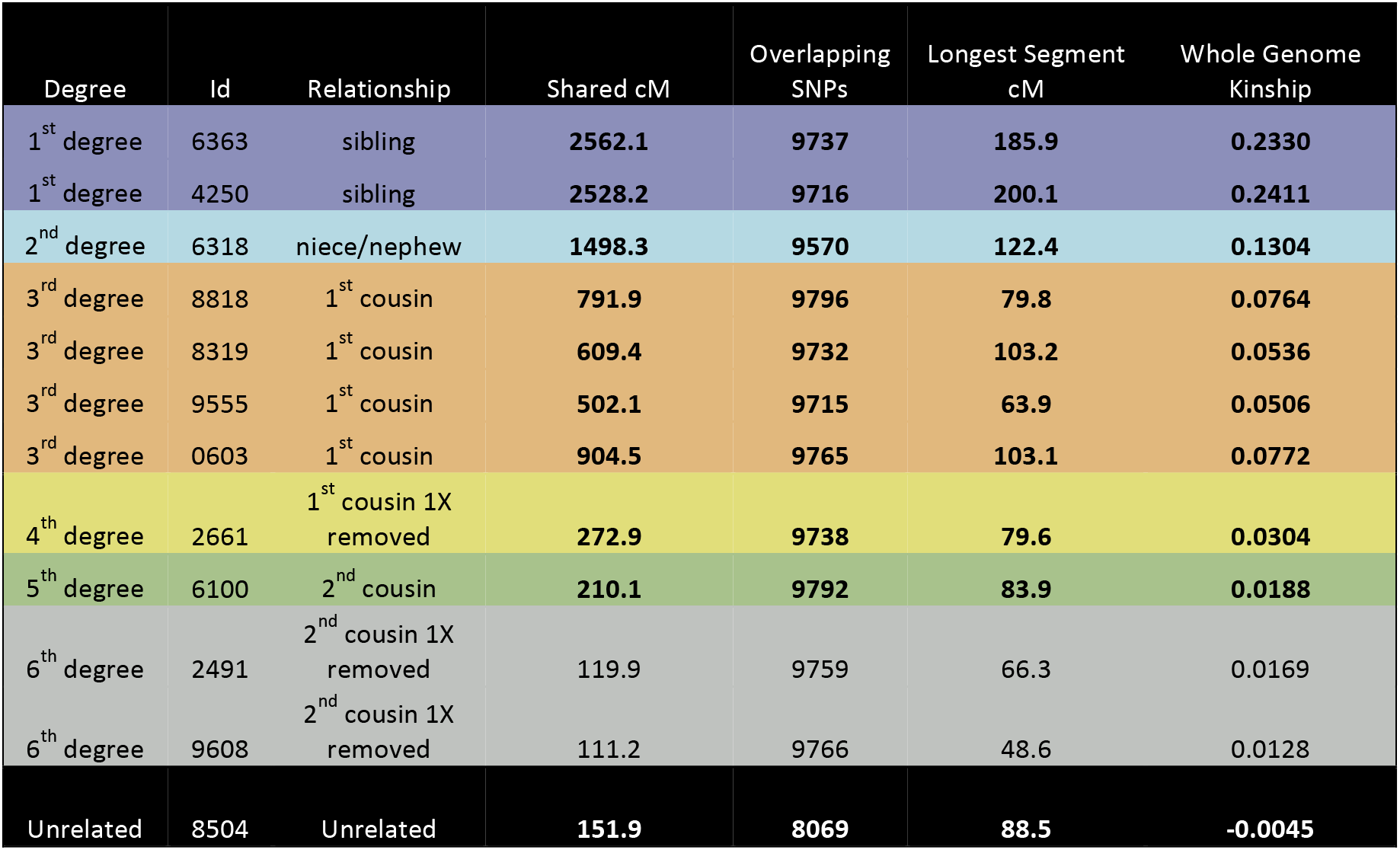
GEDmatch query results for a 10K SNP profile from sample V024 and a known pedigree using windowed and whole genome kinship algorithms. Shared cM and Longest Segment cM are calculated from windowed kinship and whole genome kinship from the standard PC-Relate algorithm. Bolded values are higher than GEDmatch Pro default thresholds. The fifth degree relationship that was detected was a false association to a synthetic profile.

The false positive profile was later identified to have many contiguous missing sections of the genome which were incorrectly being identified as extensions to segments of kinship. Later revisions to the algorithm (deployed to GEDmatch Pro in June of 2022) address this issue and the false positive is removed.

### 4.2. Windowed Kinship vs Whole Genome Kinship

The windowed kinship algorithm is a modification of the PC-AiR and PC-Relate tools for inference of genetic relatedness that use a whole genome kinship approach. Performance of the 10K SNP multiplex with the whole genome kinship approach achieved detection of associations out to the third and was improved for more distant relationships by implementing the windowed kinship approach. For close relatives, the associations detected by whole genome kinship and windowed kinship are the same as there are many overlapping SNPs across the entire genome. For more distant relatives such as second cousins once removed, the number of matching genotypes at the 10K SNP loci for related sample pairs and for unrelated sample pairs is similar. In the samples shown in Fig. S9, for the second cousin match V024 and 9608 there are 3,956 fully matching genotypes. For the unrelated pair there are 3,956 fully matching genotypes. Related and unrelated pairs can therefore produce similar whole genome kinship values. However, in the related pair of samples V024 and 9608 (see Fig. S8 for pedigree), there are distinct segments of kinship which are not the case in the V024 and unrelated sample. Given that true relatives have regions of the same SNP allele calls contiguously on a chromosome rather than randomly distributed throughout the genome, there is a much higher chance of being related if they share SNPs in contiguous blocks (even if two samples have the same number of overlapping SNPs). This concept is the same as that for segment matching, *i.e*., it is much more likely to find a segment of shared relationship than for SNPs to randomly match through the genome in true relatives.

To compare general performance of windowed kinship versus the whole genome kinship method on SNPs of the 10K set, the GEDmatch test sample set was used that contains profiles of putative relatives based on standard segment matching (see Section 3.1). Only pairs of profiles with >9,000 mutually called loci were used so that aggregate statistics were comparable. ROC curves were generated for whole genome kinship and windowed kinship methods (Fig. 4). Thresholds for windowed kinship were tested between zero and 3300 cM in steps of five for total shared cM, and between zero and 50 cM in steps of two for longest shared cM segment. Thresholds for whole genome kinship were tested between zero and 0.5 in steps of 0.01. A ROC curve that hugs the upper left-hand corner of the graph represents ability to resolve relationship classes and was observed for first through third degrees when either the windowed kinship approach or the whole genome kinship approach was used. These data indicate that 100% sensitivity and specificity can be achieved from either method in this range of relatedness. At fourth and fifth degree relationships, differences in sensitivity and specificity were observed between the two algorithms. At fifth degree in particular, the discriminatory power of the windowed kinship approach was higher than with the whole genome kinship method. From a practical perspective, given a database of 1.5 million samples and cM thresholds that support approximately 50% sensitivity for fifth degree relatives, seven false associations would be expected using windowed kinship as compared to more than 2,000 false associations using the whole genome kinship method alone.

**Fig. 4:**
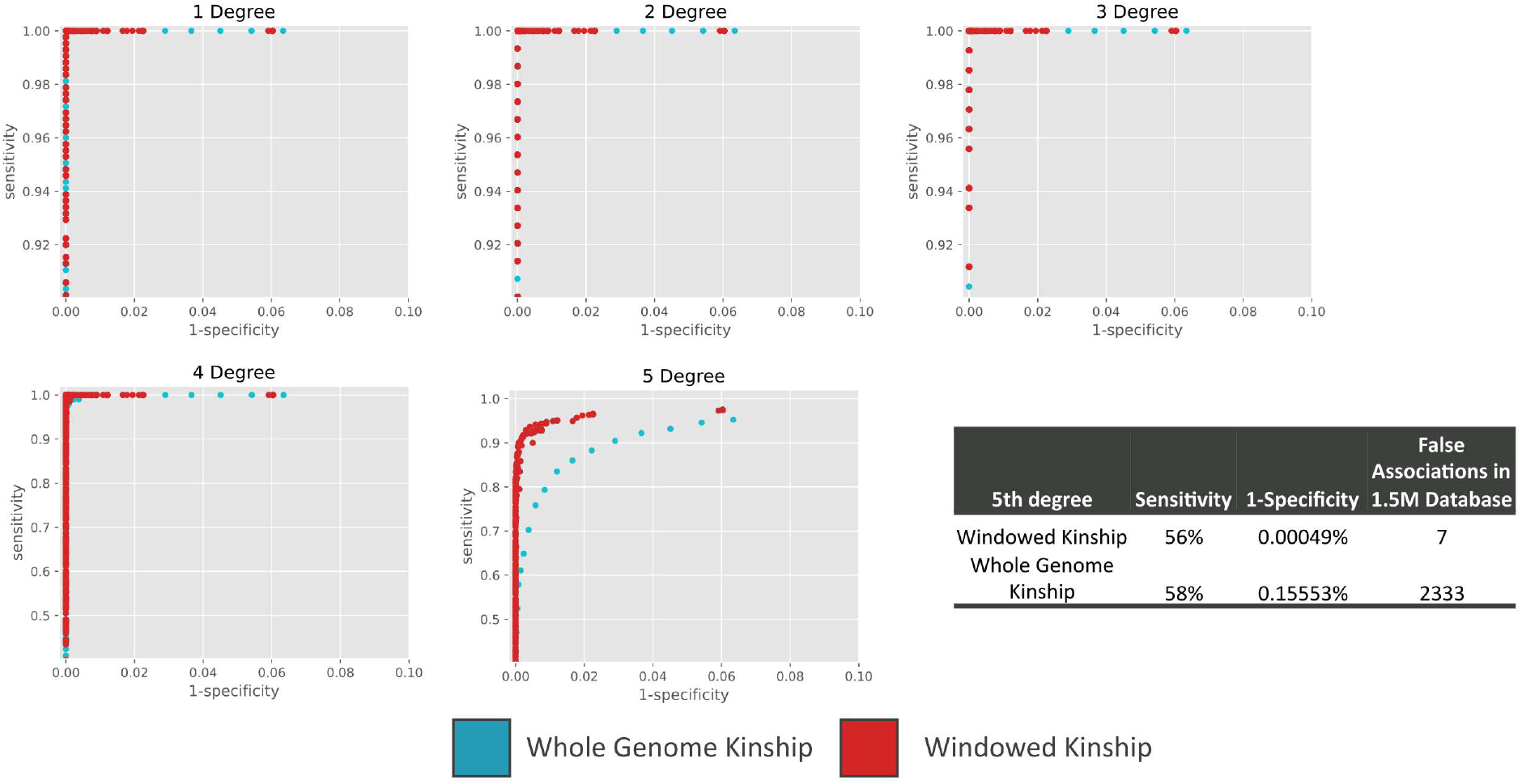
ROC curves for whole genome kinship (blue squares) vs windowed kinship (red squares) methods on a test sample set in GEDmatch comprised of the 10K SNP set. For fourth and fifth degree relationships, windowed kinship significantly improved sensitivity and specificity. When thresholds for shared total cM (for the windowed kinship method) and kinship coefficient (for the whole genome method) are set to give approximately the same sensitivity for fifth degree relationships, more false associations are detected with the whole genome kinship method (>2300); see table inset.

### 4.3. Estimated Shared cM from Windowed Kinship vs from GEDmatch Segment Matching

Segment matching algorithms used by DTC genetic genealogy companies output an aggregate metric of total shared cM. Since this metric is widely used, there are several tertiary tools that can be used to interrogate genetic kinship associations by looking at shared cM values. For example, the Shared cM Project provides an aggregate of shared cM values across many degrees of relationships, facilitating determination of what types of relationships correspond to which ranges of shared cM values.^4^ Even though the mechanism of windowed kinship is not the same as segment matching, the windowed method can provide matching segments across the genome, and output total shared cM as a kinship metric.

Estimates of shared cM from segment matching and windowed kinship were compared. One difficulty with such an evaluation is that different genetic genealogy companies use different cM maps which can lead to divergent measurements (see Fig. S11). The windowed kinship implementation in GEDmatch Pro uses newer maps from Bherer *et al*. [17] that have a total sex-averaged cM across the autosomes of 3,342 cM while GEDmatch segment matching uses an older cM map that has a total of 3,586 cM. Thus, *on average* the estimates from windowed kinship are expected to be approximately 7% lower than those with the GEDmatch segment approach. However, since the differences are unevenly distributed, they can be higher or lower depending on the shared segments between two samples.

One other issue with comparing the shared cM metric is that GEDmatch only considers half-matches in its one-to-many tool^5^. When there is at least one allele in common at a single biallelic locus, then half-matching considers that as a match between samples. For example, if there is a locus with a heterozygous call in one sample and a homozygous call in another sample, then that locus is considered a half-match since either allele from the heterozygote can match to the homozygote. For a full match, each sample must be heterozygotic or must be homozygous for the same allele to be considered matching. As relationships get more distant, it is more likely that segments of shared kinship will be comprised of more half-matches than full matches, which is sufficient as a first pass when conducting database searching. However, a “self-match”, *i.e*., two samples from the same individual, has a maximum cM value of 3,586, same as a first degree relative. Since windowed kinship considers full matching, a self-match is represented by a number closer to 6,642 cM. Therefore, samples with values from windowed kinship greater than 3,342 cM will not be the same as what is reported from segment matching in GEDmatch. To control for this effect in this study, all GEDmatch test pairs with an estimated shared cM of > 3,600 by the windowed kinship method were not considered. (thus, numbers of first degree sample pairs differ between Fig. 5 and Table S2)

**Fig. 5:**
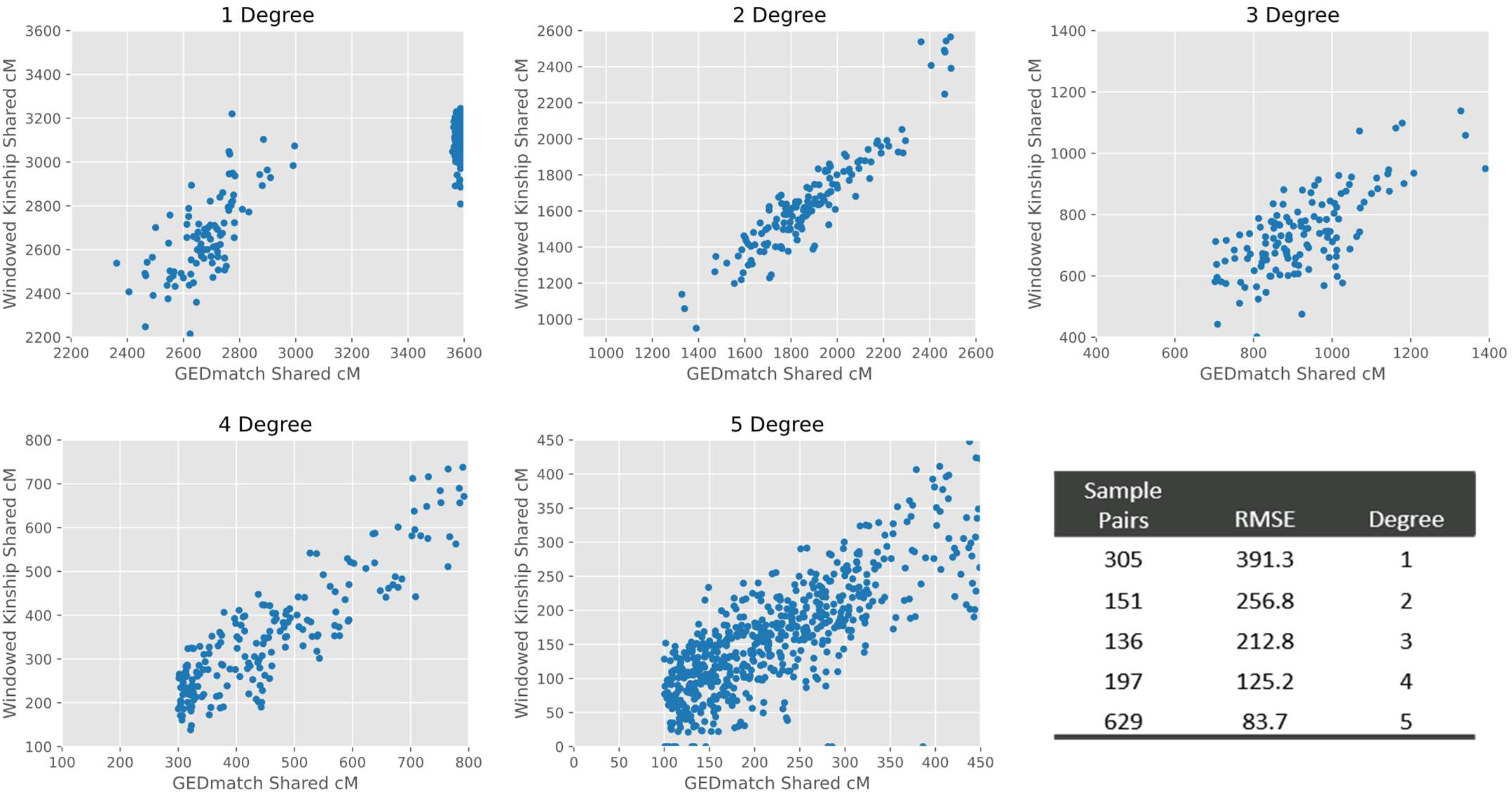
Comparison of estimated shared cM from the GEDmatch segment approach (x-axis) and the windowed kinship approach (y-axis) for 1,420 sample pairs. The inset table shows the root mean squared error (RMSE) of the windowed shared cM vs the GEDmatch shared cM estimates for the first through fifth degrees. These two approaches use different reference maps to estimate cM distances such that the overall estimated total cM in the GEDmatch segment approach differs from windowed kinship on average by ^~^7%. In general, observed estimations from the windowed kinship approach were slightly lower than from segment matching. 208first degree relationships with estimated shared cM close to the maximum possible value from GEDmatch (3,600 cM) showed lower values from windowed kinship, though still within range of a first degree relationship.

With these caveats in mind, concordance and differences between cM estimates from these two methods were compared. As shown in Fig. 5, the estimates of total shared cM between the windowed kinship approach and GEDmatch segment matching were similar, although variability between them was observed. Interestingly, there are many GEDmatch first degree hits with values close to the maximum possible shared cM values that have a wider spread for windowed kinship. The windowed kinship estimates fall within the first degree shared cM ranges. These data indicate that differences in the total shared cM values may be observed for close relatives compared to values generated with the GEDmatch segment approach. This difference may be due to segment matching less aggressively filtering segments than the windowed kinship approach thus is not an impediment to conducting FGG.

### 4.4. Windowed Kinship Performance with Forensic Case-Type Samples

#### 4.4.1. Windowed Kinship Performance with Partial Profiles (<10K Kinship Informative SNPs)

Some loci in targeted assays of forensic samples or unidentified human remains may not be detected (*e.g*., data below an analytical threshold or no data detected) such that partial profiles are generated due to DNA degradation, damage and/or PCR inhibition. To assess performance of the kinship algorithms for samples with different levels of missing loci, the GEDmatch truth set described in Section 3.1 was used. The GEDmatch test set used standard segment matching on microarray data to locate relatives and then the test SNPs were filtered to the 10K SNP set and used for kinship inference. In this evaluation, from the set of 10K SNPs, random subsets of loci were selected and marked as missing from the input profiles of the GEDmatch test set. Between 2000 and 8000 loci were removed in this fashion and evaluated, equivalent to 80-20% SNP locus call rates.

ROC curves were used to evaluate these data as different levels of missing loci can be recognized when kinship is estimated. For example, if the specificity in estimating relationship degree is reduced when a certain number of the 10K loci are untyped, then kinship thresholds can be adjusted dynamically to account for it. As shown in Fig. 6, at lower levels of missing loci and out to fourth degree relationships the ROC curves were sharply upper and leftward, indicating high sensitivity and specificity. Out to fourth degree, SNP locus call rates greater than 60-80% generated similar results to those from full 10K SNP profiles. For close relationships (first to third), performance was maintained down to a 40% SNP locus call rate. Although similar performance can be achieved with large numbers of missing loci, the kinship thresholds necessary to achieve that performance can differ. Thus, it is important to use different thresholds based on how many SNPs are shared between samples, *e.g*., if a pair of samples has 6,000 SNPs typed in common, a higher windowed kinship threshold can be used than for a pair of samples with 9,000 overlapping SNPs (see Table S4 for thresholds set for windowed kinship in the GEDmatch Pro implementation).

**Fig. 6:**
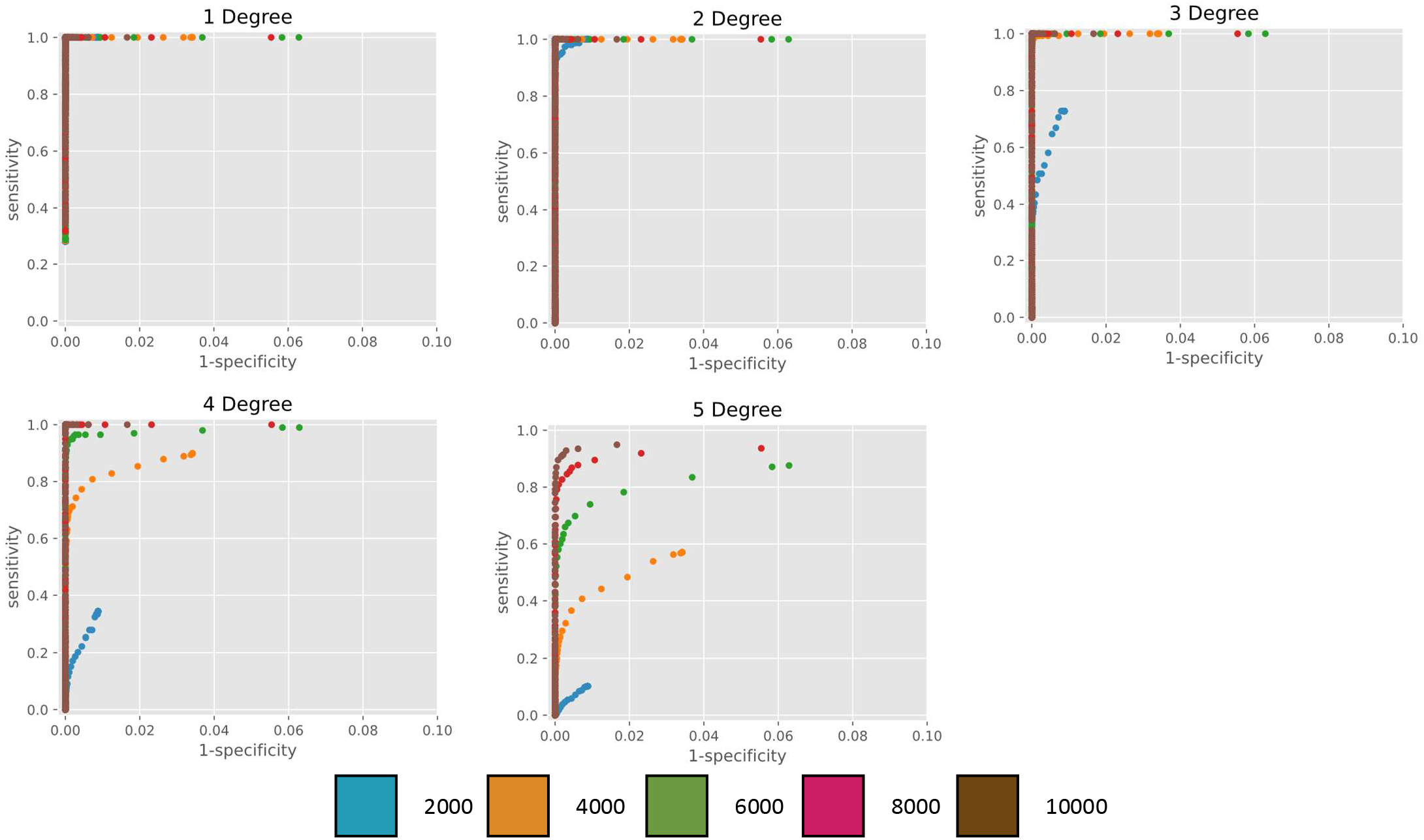
Performance of windowed kinship in GEDmatch test set with call rates between 20-100% for the 10K SNP set. Data are plotted for five locus call rates as follows: 20% (2K), 40% (4K), 60% (6K), 80% (8K) and 100% (10K). Overall, performance for first, second and third degrees was observed to be steadily maintained when 80%, 60% or 40% of the 10K SNPs were typed. For fourth degree, 80% of the 10K SNPs were observed to give comparable performance to the full 10K set, and 60% was sufficient to make some kinship analyses (approximately 60% sensitivity vs 100% sensitivity for the full profile at the same specificity).

##### 4.4.1.1. Windowed Kinship Performance using Partial SNP Allele Call Rates

Total heterozygosity at SNP loci in a human DNA sample and quantitative balance between heterozygous alleles can be used as quality metrics for SNP genotyping and particularly for assessment of profiles from challenging samples, including those in forensic casework [21]. These metrics can indicate the likelihood that only one of the sister alleles in a true heterozygote were detected and may be called as homozygous. As sample quality degrades and input DNA template is reduced, certainty in homozygous SNP calling can be affected. Forensic genetics casework employs methods and tools to assist in this regard, such as use of stochastic thresholds [22][23]. For the windowed kinship algorithm, whether similar threshold(s) are necessary to disqualify SNP data outright from proceeding with FGG or whether the algorithm was robust to some missing alleles was investigated.

To evaluate how loss of sister alleles affects windowed kinship performance, the GEDmatch test set was used. As with the previous evaluations, the truth set was generated from segment matching on whole microarray profiles, and the samples were filtered to the 10K SNP set. Different percentages of heterozygous loci were changed to homozygous reference (ref) or alternative (alt) calls. Ref calls generally refer to the more prevalent allele in a reference population while alt calls refer to the less prevalent, or “minor” allele. For the SNP locus rs6690515 as an example, a G is considered “ref” while A is considered “alt”. Converting a G/A call to a G/G call, changes a heterozygote to a homozygous ref call. The ref allele is represented as 0 and the alt allele as 1 when the actual nucleotide is not germane.

An example of the simulation strategy used in this study is as follows: Consider a simulation of 5% of sister alleles at heterozygotes among the 10K SNP set. The transition probabilities of the genotypes from the original profile are shown in Table 2. The transition table provides the percentage of a heterozygous locus modified to a homozygous call in the test case simulations of allele non-detection. For example, if an input sample has a locus with a starting genotype of 0/0, the test profile will also have a genotype of 0/0 since the probability that 0/0 transitions to 0/0 is 100%. However, if the starting genotype is a 0/1 genotype, the chance was 95% to remain 0/1 and 2.5% chance to become 1/1 or 0/0, indicating non detection of a sister allele. Essentially, this emulates cases where the second allele in a heterozygote is below the analytical threshold and therefore, calling a heterozygous call as a homozygote erroneously.

**Table 2:** Transition probabilities for 5% lack of detection of sister alleles at heterozygous SNPs as used in simulation studies. ref allele (0), alt allele (1).

| Original Genotype | Test Genotype | Probability |
| --- | --- | --- |
| 0/0 | 0/0 | 100.0% |
| 0/1 | 0/0 | 2.5% |
| 0/1 | 0/1 | 95.0% |
| 0/1 | 1/1 | 2.5% |
| 1/1 | 1/1 | 100.0% |
| ./. | ./. | 100.0% |

Ranges of missing sister allele calls between 5 and 100% were tested. Whereas with missing loci it is trivial to determine how many are missing, it is more difficult to quantify sister allele loss in an unknown sample since it can depend on factors inherent to the sample and to the subpopulation of origin. It is likely then more illustrative to analyze performance using the default windowed kinship thresholds than all possible thresholds (see Fig. S6 for full ROC). Using the default kinship thresholds for the windowed algorithm as implemented in GEDmatch Pro (Table S3), sensitivity was observed to be maintained for first to fourth degree relationships when loss of sister allele detection was less than 10%. When 20% of sister alleles were not called, kinship performance was maintained within the first to third degrees. At a 40% loss performance was maintained within the first and second degrees and at greater sister allele loss only first degree were captured (Fig. 7). Crucially, specificity was similar across all levels of heterozygous allele call rates, indicating that the loss of sister alleles did not introduce false associations.

**Fig. 7:**
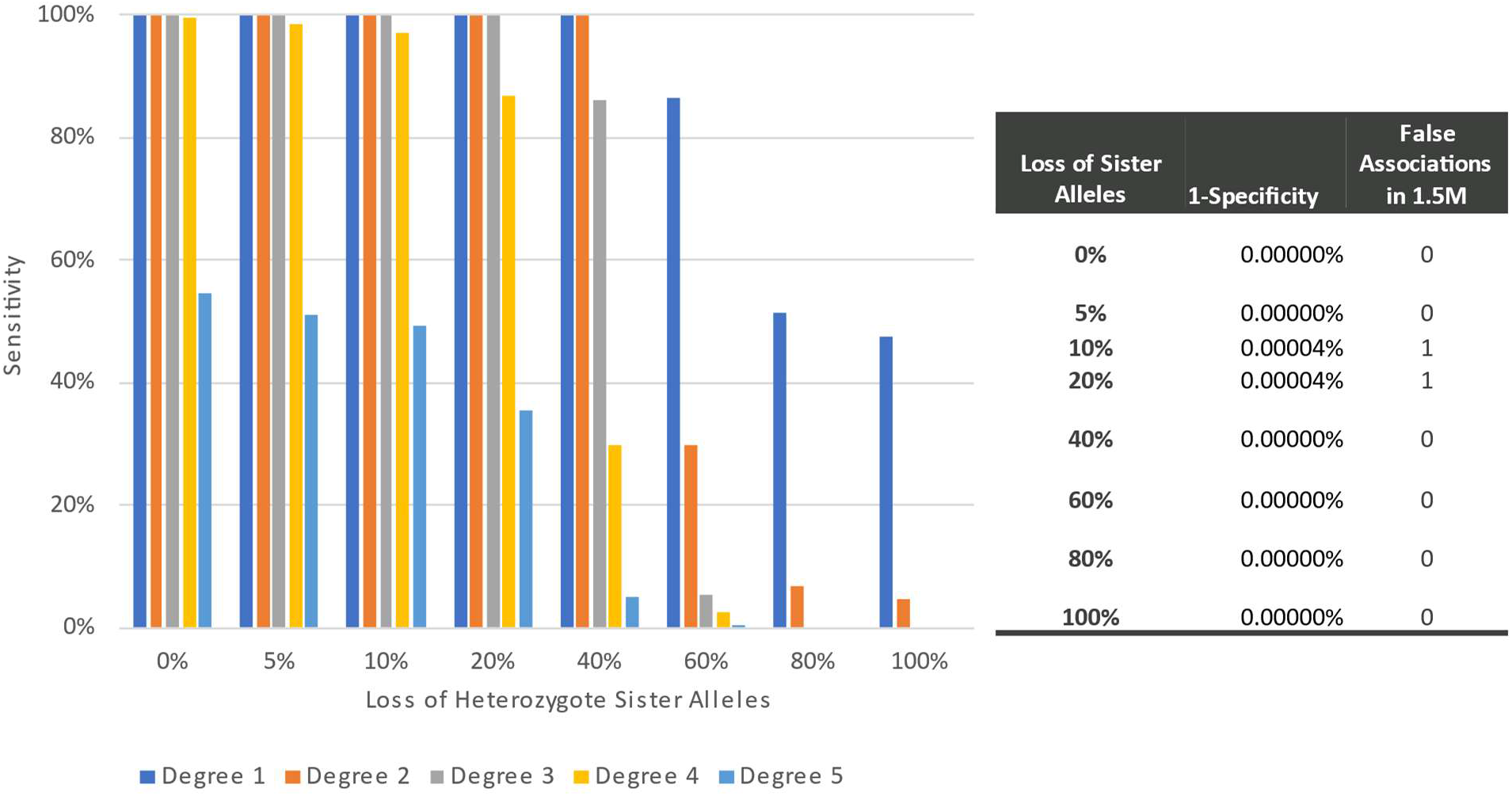
Performance of the windowed kinship method on the GEDmatch test set after simulating loss of sister alleles (between 5 and 100%) at heterozygous sites of the 10K SNP set. Kinship thresholds are based on default settings of the windowed kinship algorithm implementation in GEDmatch Pro for profiles with > 9,000 SNPs typed in common (overlapping) (i.e., 140 shared cM total, 30 cM longest segment). Simulated losses were evenly distributed between ref and alt alleles, i.e., a heterozygote in the fully typed profile had a 2.5% chance to become homozygous alt and a 2.5% chance to become homozygous ref. 1-specifitiy indicates the chance of incorrectly classifying an unrelated association as related, sensitivity indicates the percentage of total true associations found above threshold.

In addition to simulated data, a known pedigree (see Fig. S11) containing relatives extending to the fifth degree was used. The person of interest (“self”) sample (V016) was heat treated to emulate partial DNA degradation and windowed kinship metrics generated from two PCR template inputs (1 ng and 250 pg for Kintelligence library preparation for sequencing) and compared. In order to test the limits of the system, one sample was run at higher plexity (12 samples in a run) than recommended by the manufacturer and also used 250 pg input. For these empirical samples, the expected associations out to third degree passed GEDmatch Pro thresholds for the 1 ng sample and second degree for the 250 pg sample (7% heterozygosity).

As represented by total shared cM values (Fig. 8) as sister allele non-detection increases, the overall estimated shared cM value dropped. For example, in a comparison of sample V004 with sample V016 with 1 ng input they fall within the expected shared cM range for a first degree hit with 3076.6 (see Table S1). The same sample compared to a V016 sample with 250 pg input only showed a shared cM value of 1561.097, which is significantly lower than would be expected for a first degree candidate hit.

**Fig. 8:**
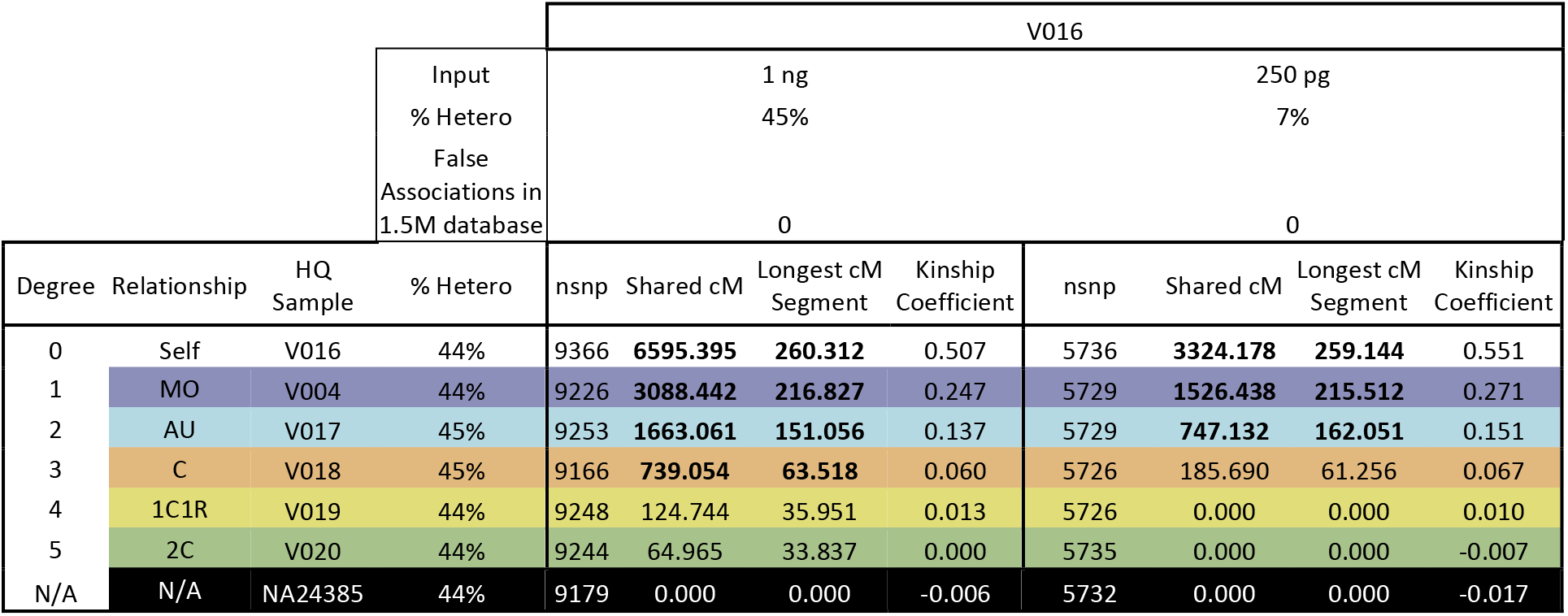
Example showing shared cM for sample V016 as a mock casework sample within a known pedigree. All samples were typed for the 10K SNP set using the ForenSeq Kintelligence Kit. V016 (self) was heat treated to emulate DNA degradation. In order to test the limits of the system, one sample was run at higher plexity (12 samples in a run) than recommended by the manufacturer and also used 250 pg input. Bolded cells indicate which values are above the thresholds used in GEDmatch Pro. Samples were also searched against full 1.5M GEDmatch database and no false associations were found above thresholds. These are currently 140 total shared cM and 37 longest cM segment for samples with more than 9000 overlapping SNPs and 180 total shared cM and 37 longest cM segment for samples with 6000 overlapping SNPs. “nsnp” indicates the total number of SNPs shared between the two samples in the pair.)

## 5. Conclusions

The windowed kinship algorithm applied to data generated from the 10K SNP multiplex supports near perfect detection of relationships extending to the third degree in a large database with a high degree of specificity even in samples with reduced locus call rates or lack of detection of sister alleles in heterozygotes. Using simulated and real GEDmatch SNP profiles, comparable performance was observed for the windowed kinship algorithm and the 10K SNP set as compared to the segment matching approach that uses hundreds of thousands of SNPs. In real degraded samples the ForenSeq Kintelligence system can identify relationships robustly out to the 3^rd^ degree. For forensic samples, the approach described herein can be considered as a powerful tool for investigative lead generation in forensic casework and unidentified human remains investigations that can be readily transferred and implemented into operational settings under an insourced or outsourced FGG SNP typing model.

## Acknowledgements

Special thanks to Bruce Budowle for editing and review assistance and to John Hayward and Wilbon Davis for reviewing sections relating to GEDmatch.

## Supplementary Material

### GEDmatch Test Set Characteristics

**Table S1:**
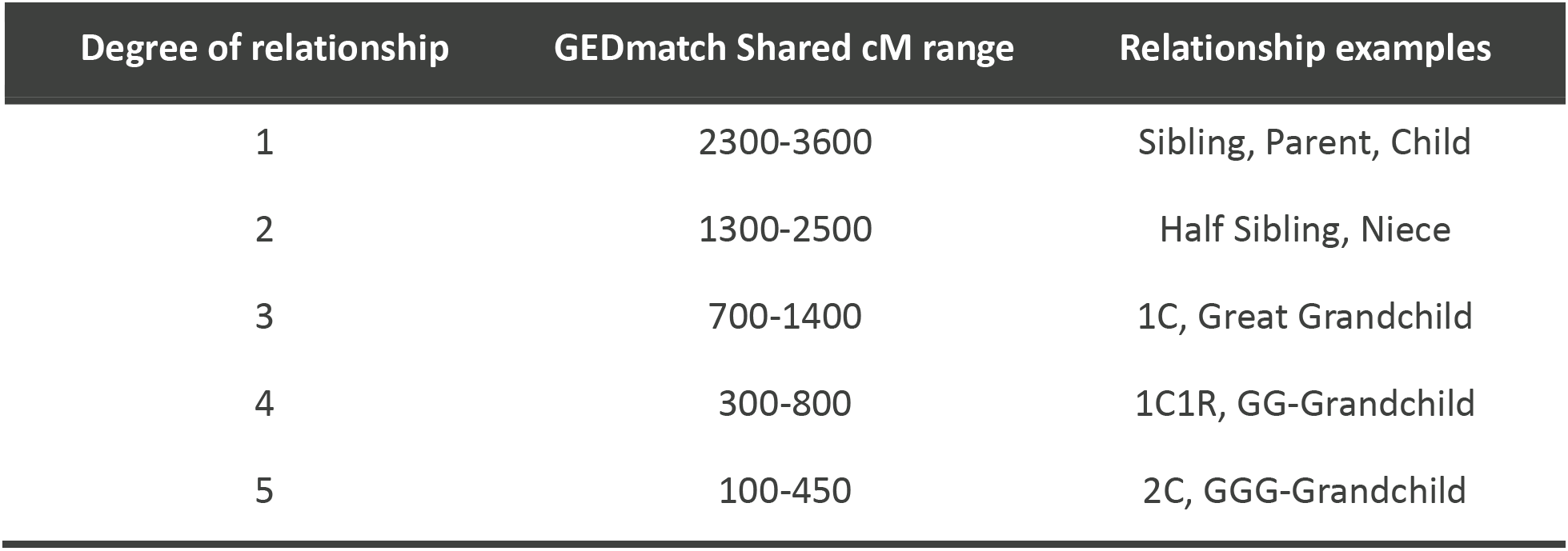
Expected shared cM ranges per degree of relationship in GEDmatch. cM ranges shown are based on DNAPainter^6^. If a pair of samples falls into more than one range (i.e., 400 shared cM overlaps with the ranges for fourth and fifth degrees) evaluation of both relationship degree possibilities may be advantageous. First cousin (1C), first cousin once removed (1C1R), second cousin (2C), great great grandchild (GG-Grandchild), great great great grandchild (GGG-Grandchild).

**Fig. S1:**
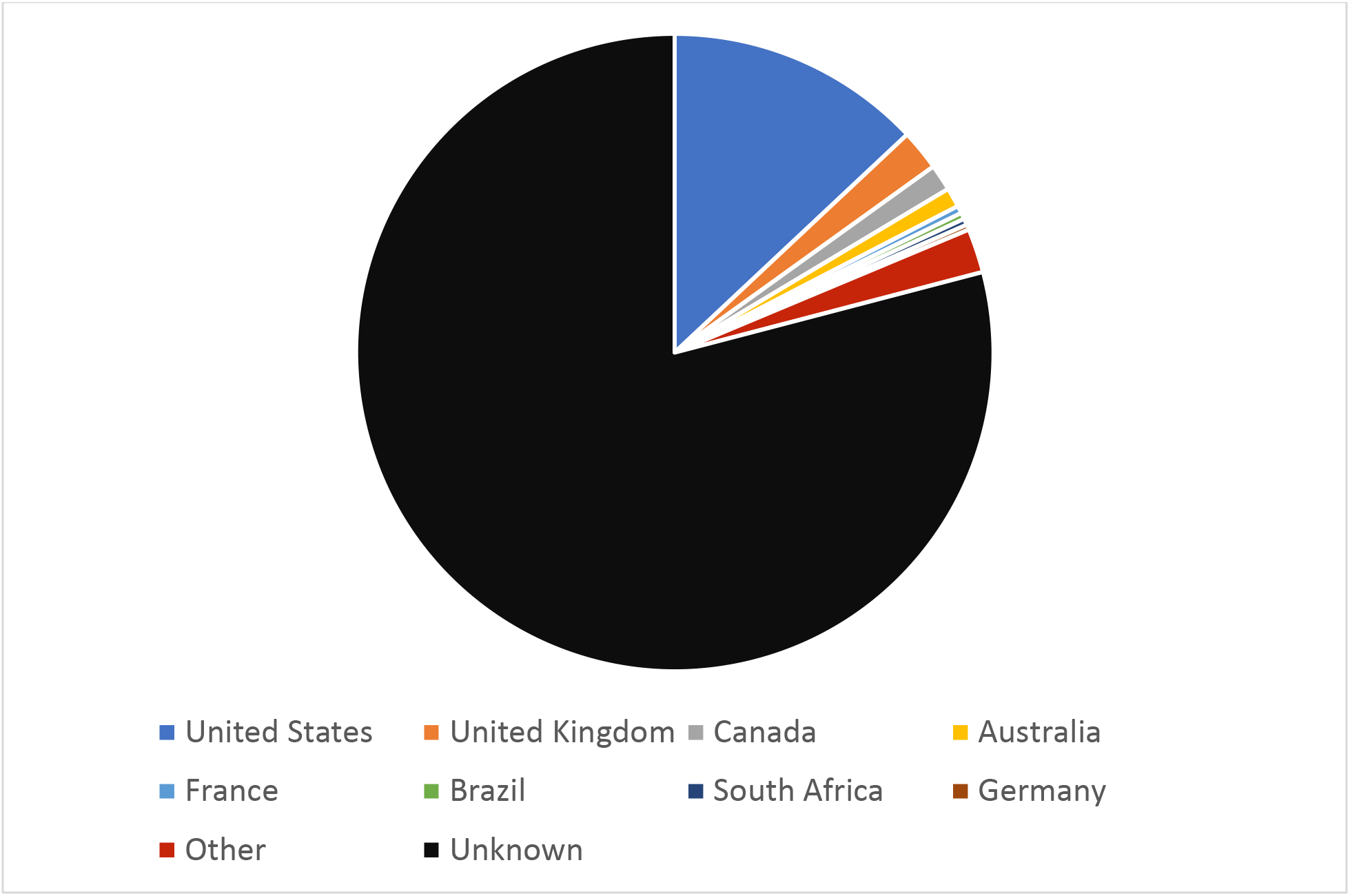
Country of origin for GEDmatch test samples based on ip-address (when available as of January 1^st^ 2022).

**Fig. S2:**
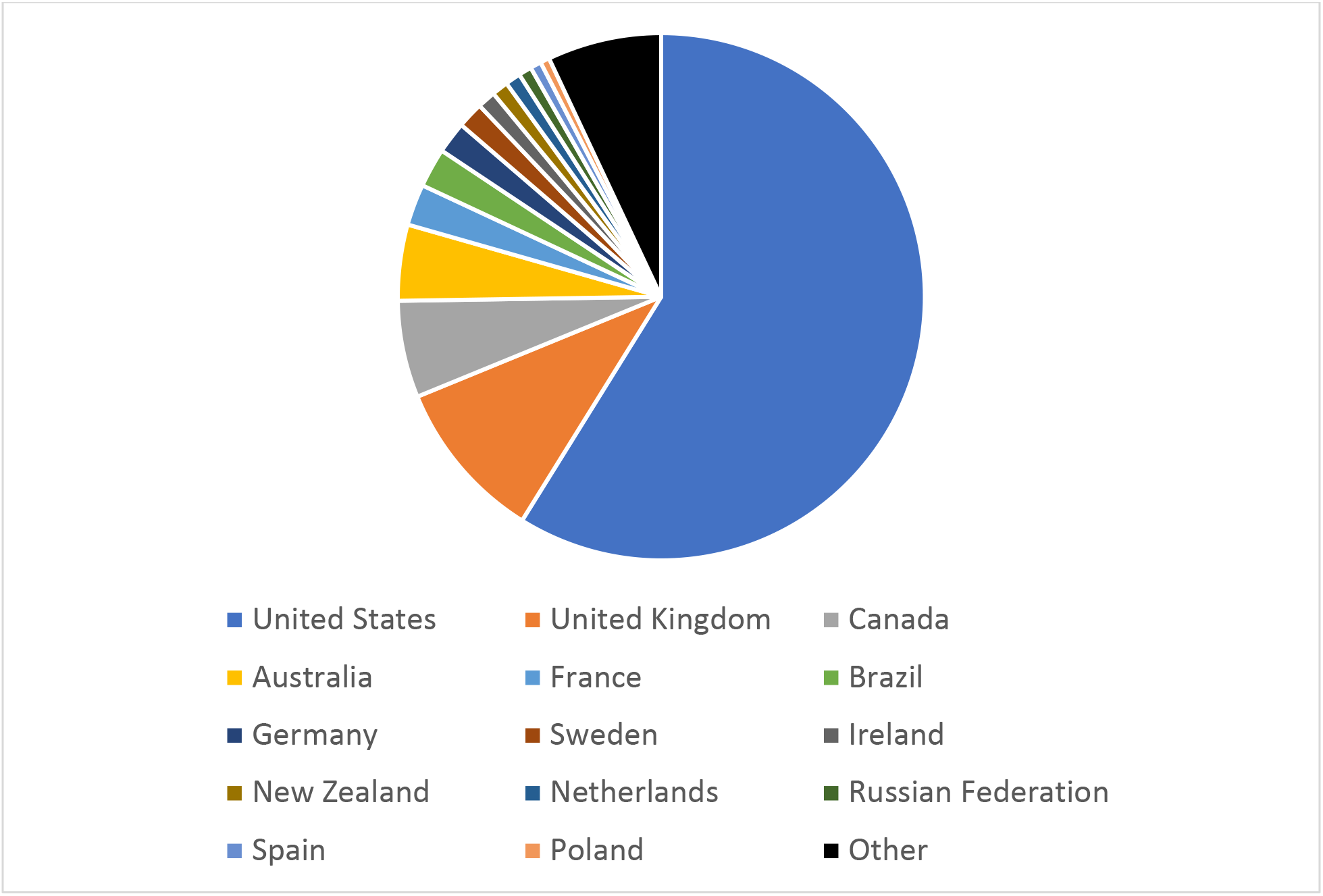
Country of origin for GEDmatch database as of January 1^t^ 2022 based on ip-address (when available)

**Table S2:** Observed shared cM ranges per degree of relationship in GEDmatch for the test set of 2,363,983 sample pairs. Pairs are limited to results where 9000 of the same loci are called for both kits.

| Degree | Shared Cm Range | Kit Pairs |
| --- | --- | --- |
| 1 | 2300-3600 | 425 |
| 2 | 1300-2500 | 151 |
| 3 | 700-1400 | 136 |
| 4 | 300-800 | 198 |
| 5 | 100-450 | 629 |
| Not Related | 0 | 2362589 |

**Fig. S3:**
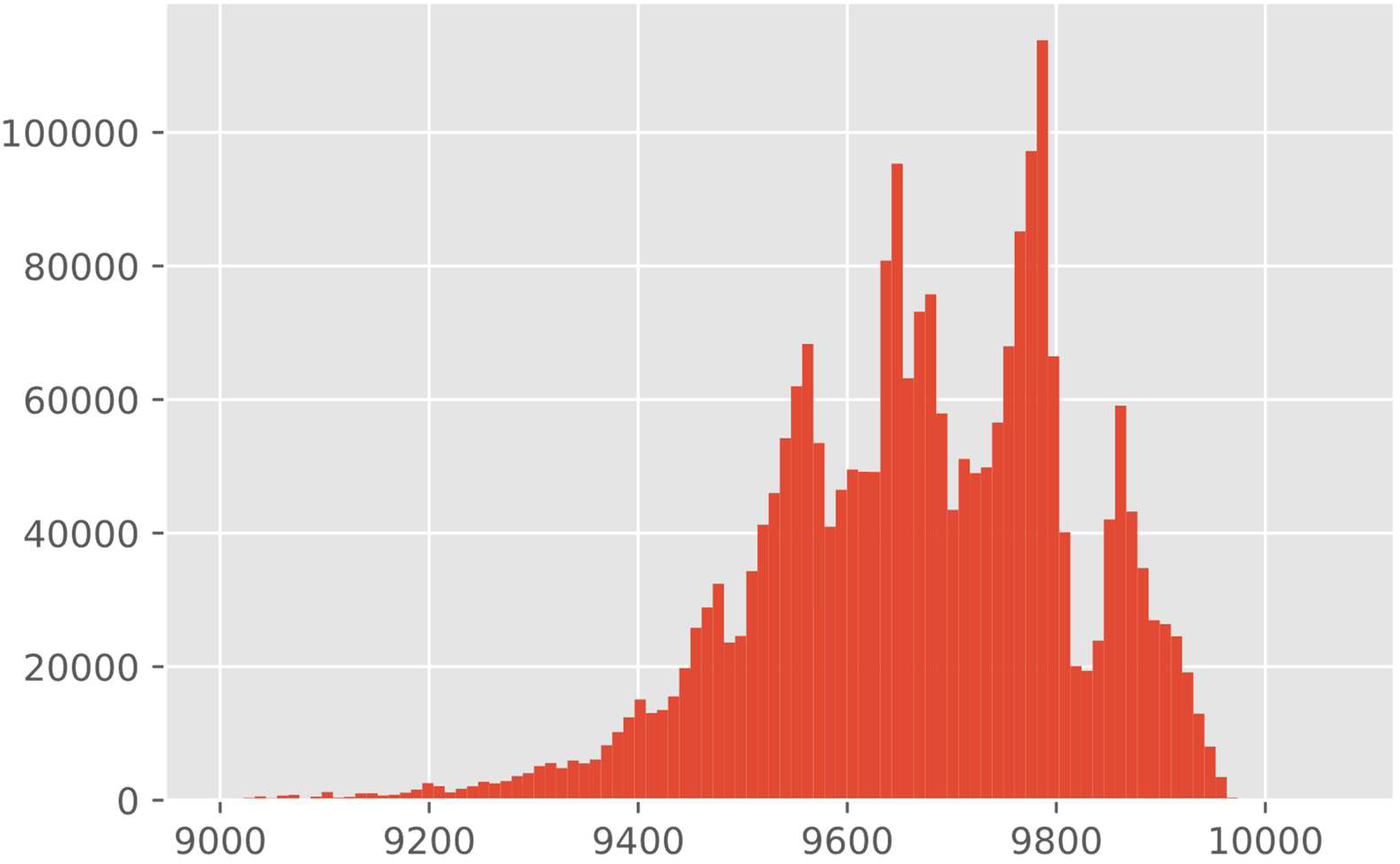
GEDmatch test set. Overlapping passing SNPs for pairs of samples from 10K SNP multiplex.

### SNP Multiplex Design

**Table S3:** gnomAD population frequencies used during selection.

| gnomAD Population | Description |
| --- | --- |
| afr | African-American/African ancestry |
| ami | Amish ancestry |
| amr | Latino ancestry |
| asj | Ashkenazi Jewish ancestry |
| eas | East Asian ancestry |
| fin | Finnish ancestry |
| nfe | Non-Finnish European ancestry |
| oth | Other ancestry |
| sas | South Asian ancestry |

**Fig. S4:**
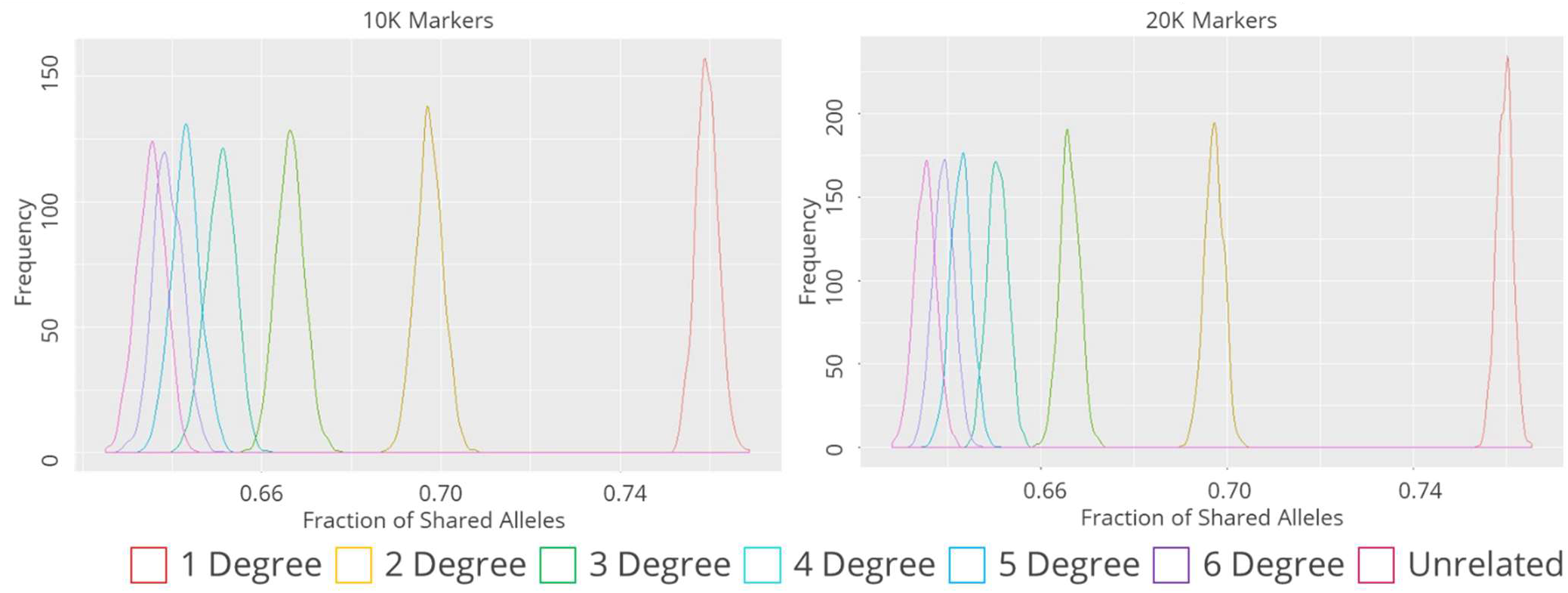
Comparison of shared allele fractions between 10K (left) and 20K (right) SNP multiplexes from ped-sim simulations across kinship relationships from first to sixth degree and unrelated. 1,000 sample pairs were generated per degree of relationship.

**Fig. S5:**
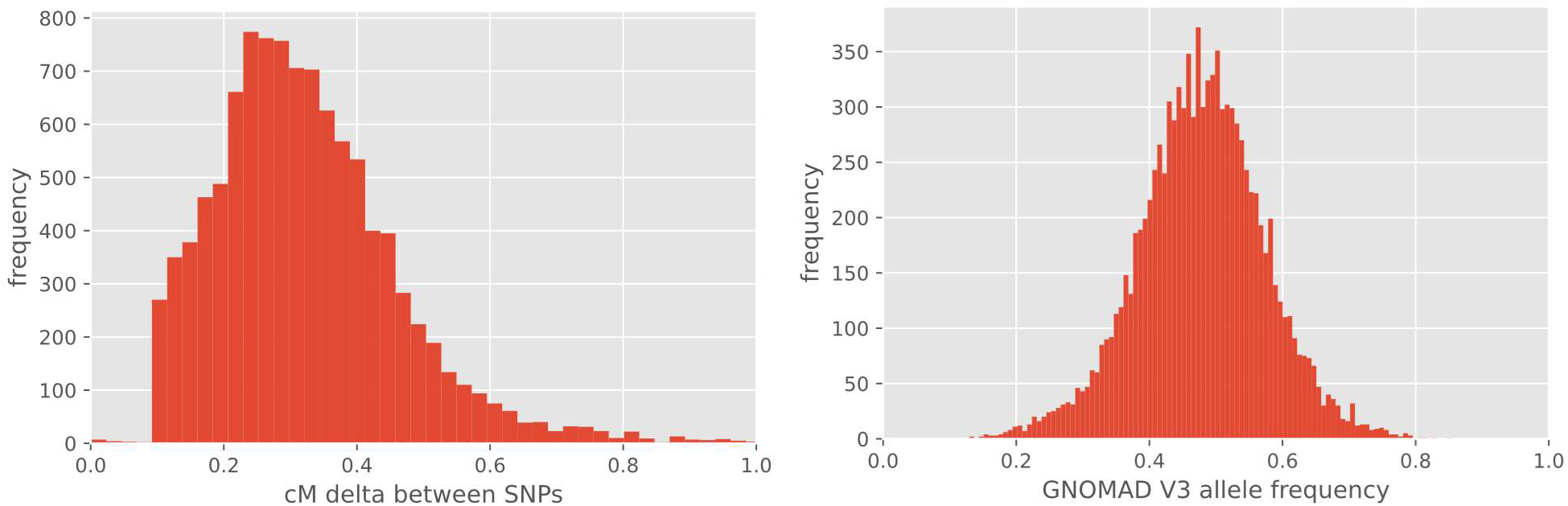
SNP characteristics for of the 10K SNP set (ForenSeq Kintelligence). The left chart displays a histogram of the cM distances between loci; the right chart displays a histogram of the gnomAD allele frequency for the SNPs in the multiplex. A minimum of 0.1 cM was required for the kinship SNPs in the 10K multiplex; loci shown here that are below that value are SNPs informative for biogeographical ancestry, phenotype estimations or identity informative SNPs from the ForenSeq™ DNA Signature Prep Kit.

### GEDmatch Pro Kinship Statistic Thresholds

**Table S4:** Windowed Kinship algorithm default thresholds as implemented in GEDmatch Pro and their influence on sensitivity of detection in first through fifth order relationships. Threshold values are based on the estimated false association (FA) rate in a search of the entire database.

| Overlapping SNPs | Shared cM Total | Longest Segment cM | Sensitivity |  |  |  |  | 1-Specificity | FAs in 1.5M Database |
| --- | --- | --- | --- | --- | --- | --- | --- | --- | --- |
|  |  |  | 1st | 2nd | 3rd | 4th | 5th |  |  |
| 9000 | 140 | 30 | 100% | 100% | 100% | 99% | 55% | 0 | 0 |
| 8000 | 150 | 30 | 100% | 100% | 100% | 99% | 41% | 0 | 0 |
| 6000 | 180 | 30 | 100% | 100% | 100% | 66% | 17% | 0 | 0 |

### Lack of Detection of Sister Alleles

**Fig. S6:**
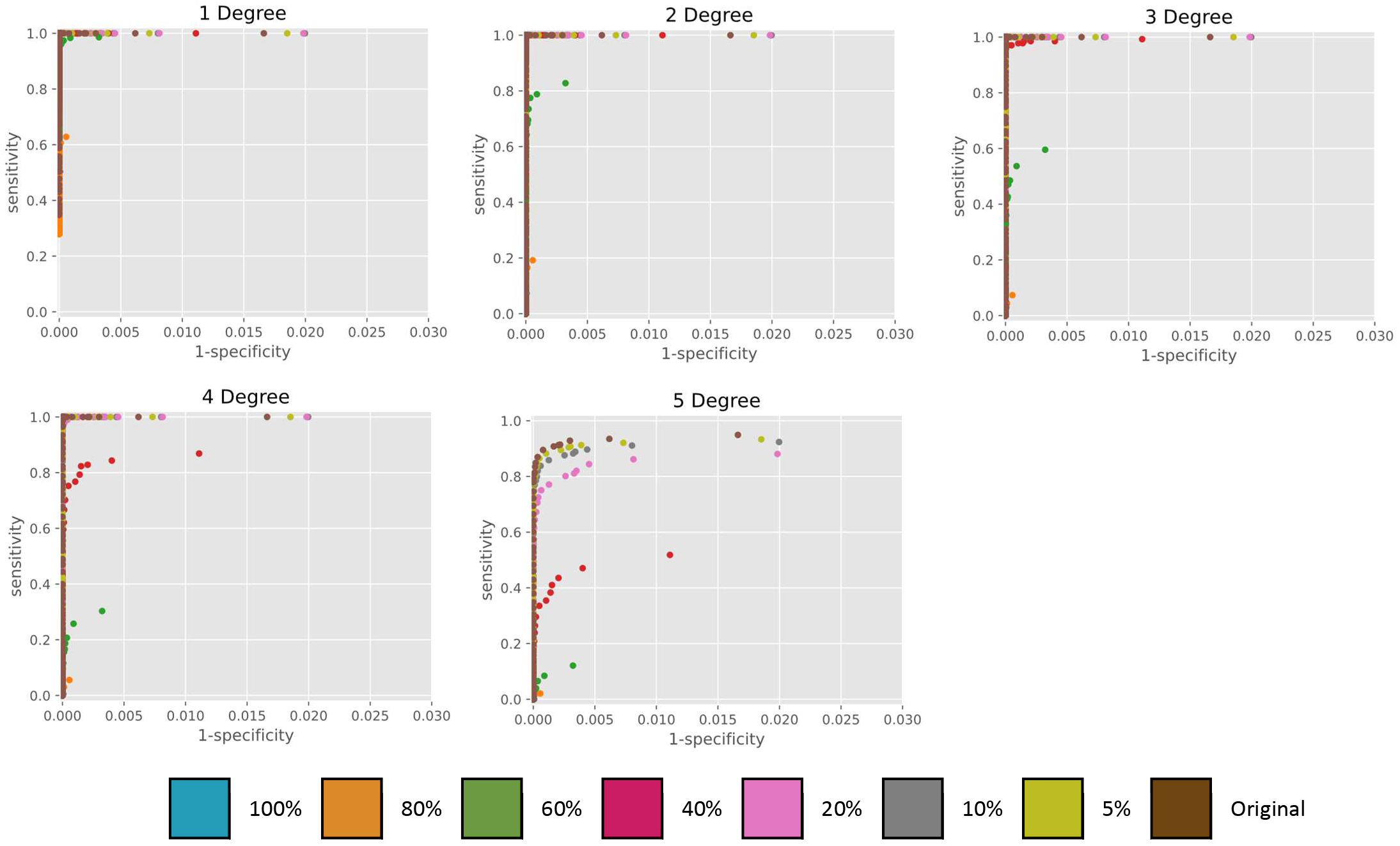
ROC curves based on GEDmatch test set using simulated losses of heterozygote sister alleles between 5 and 100%.

### Ped-sim Simulation

**Fig. S7:**
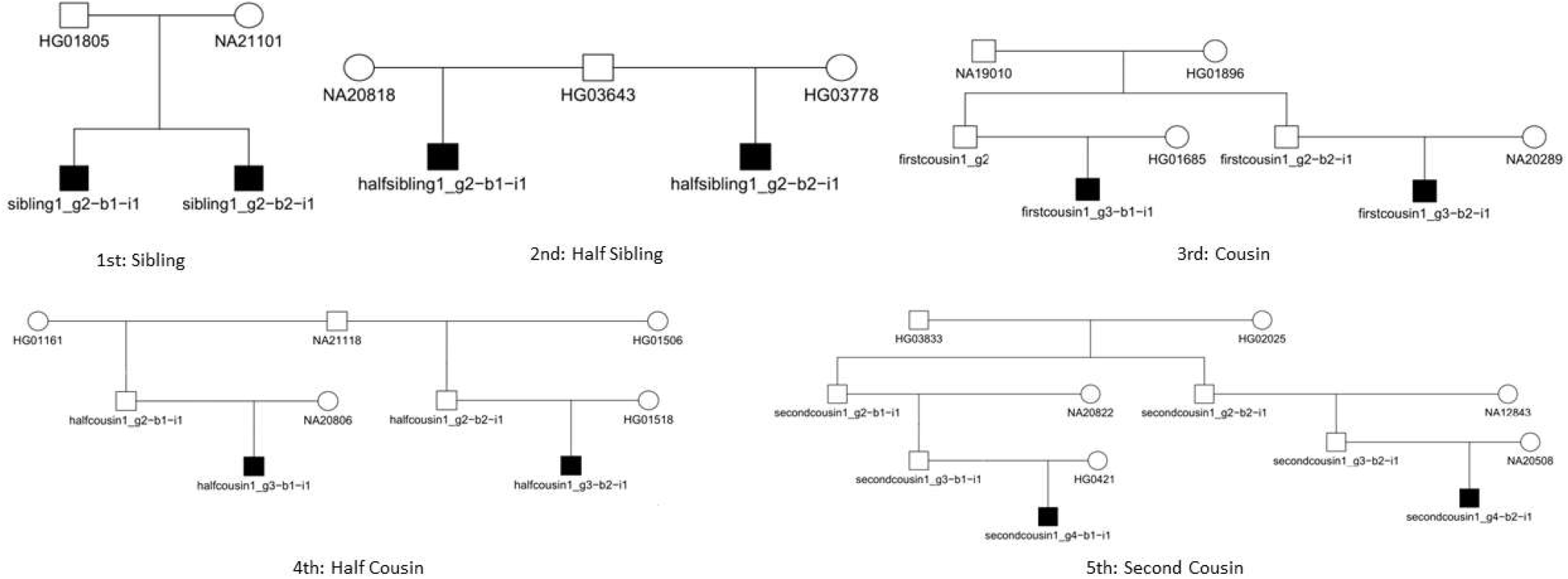
Example pedigree for first through fifth relationship degrees using 1,842 founders from 1000 Genome Project samples. An example of each degree of a single pedigree simulated by ped-sim is shown. The biological sex of the founding samples was ignored, and sex averaged linkage maps were used in the simulations. Only the darkened samples are output by ped-sim and used in evaluation scripts for this study. Pairs of samples from the same pedigree were considered true relatives, pairs of samples across pedigrees are considered unrelated.

### Known Kinship SNP Pedigrees

**Fig. S8:**
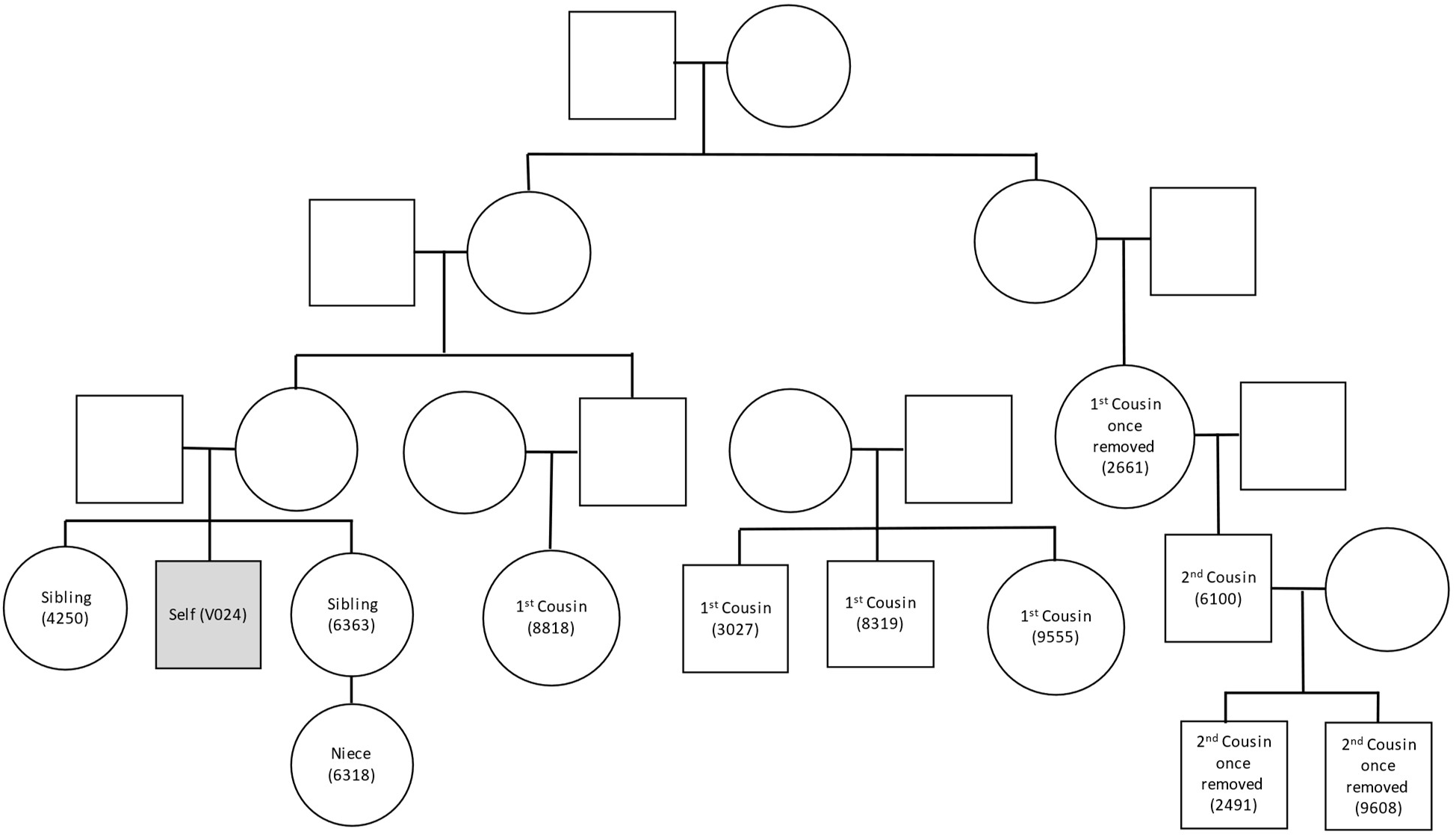
Known extended pedigree from GEDmatch. Sample V024 was assigned as the person of interest and typed for the 10K SNP set using the ForenSeq™ Kintelligence kit; all other sample profiles are from microarray typing.

**Fig. S9:**
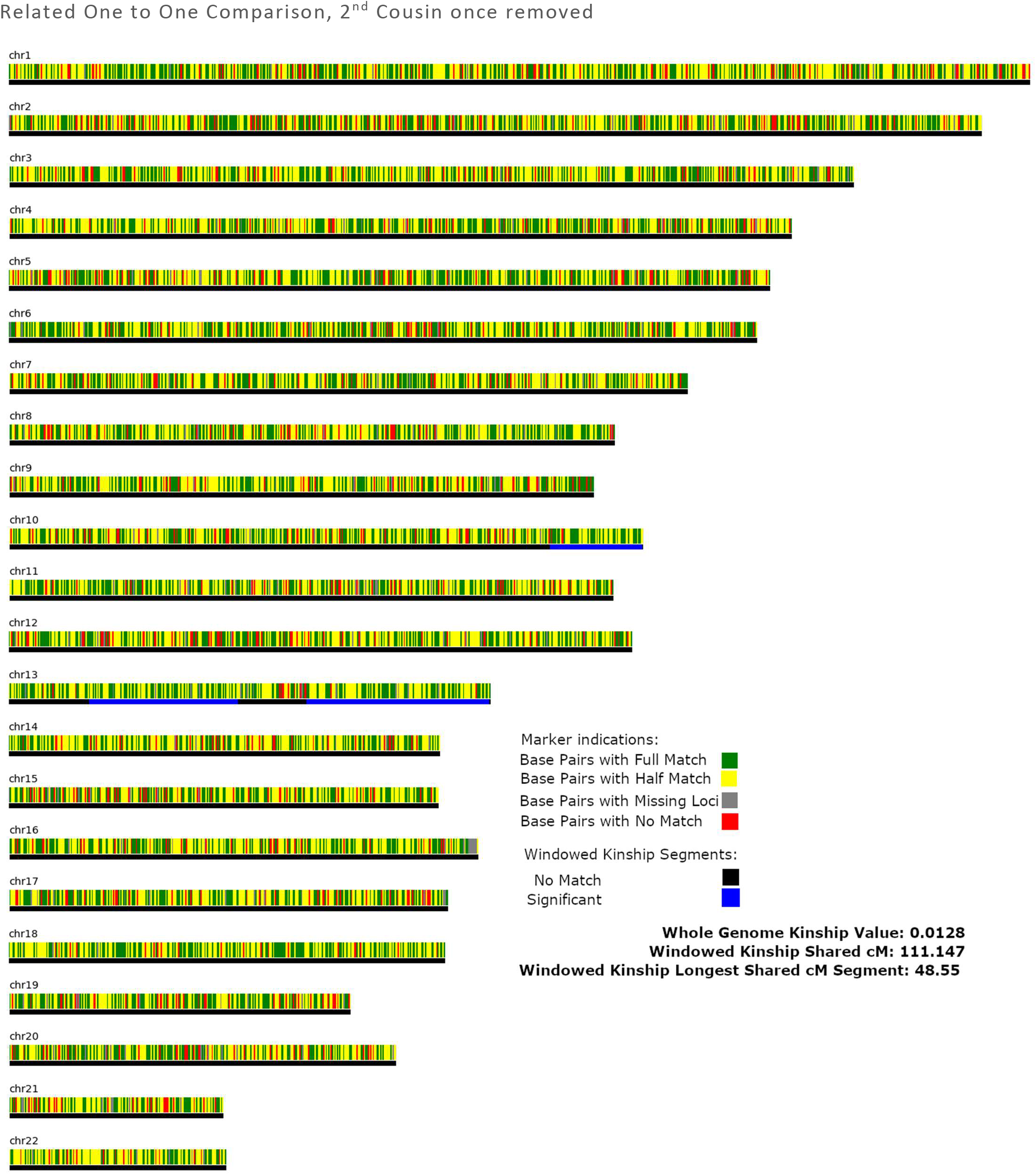
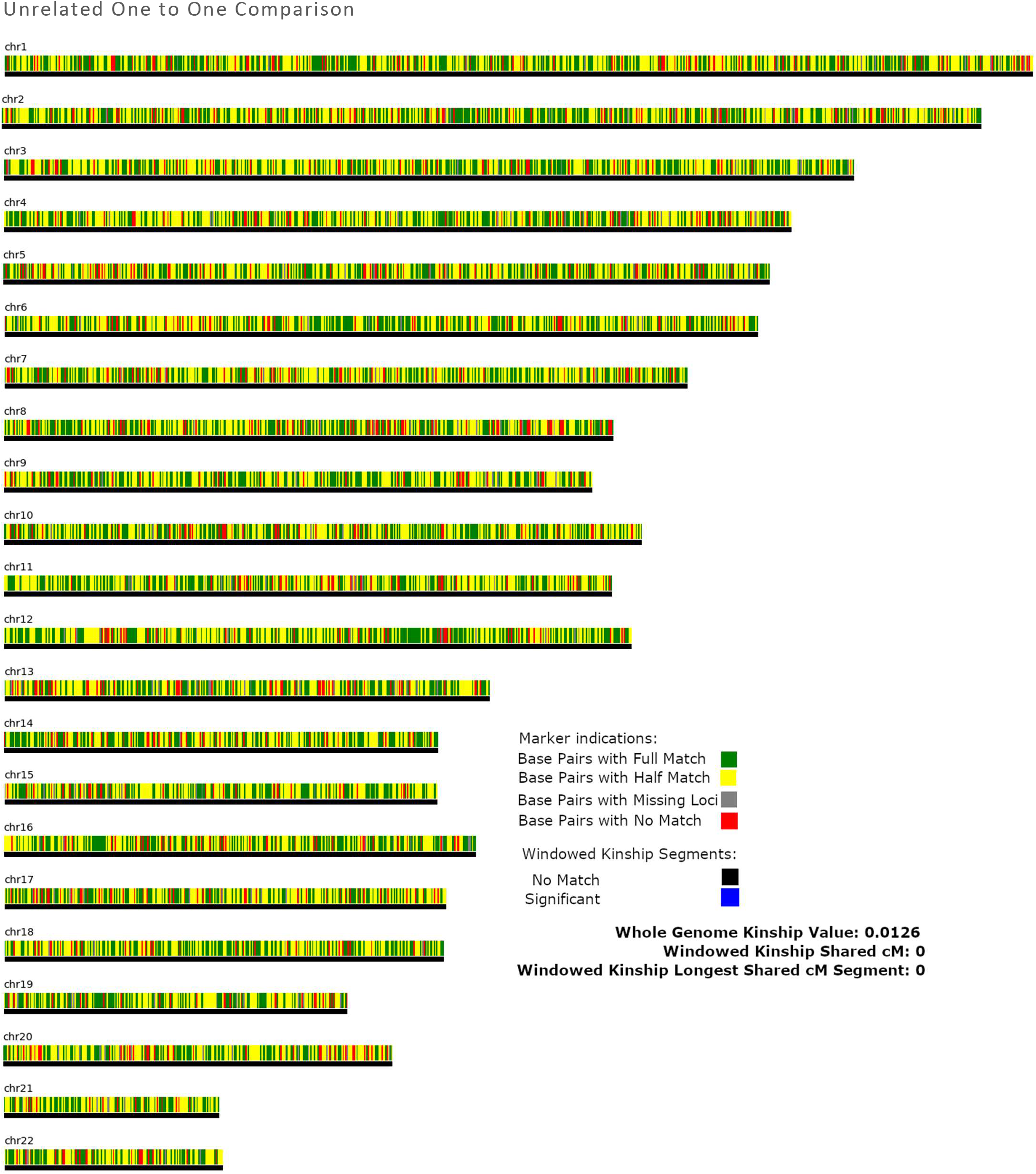
Visual display of matching SNPs across genome. The upper panel show sample of interest V024 as compared to sample 9608 a 2^nd^ cousin once removed from a known pedigree. The lower panel shows the sample of interest V024 as compared to an unrelated sample. The sample of interest was typed using the 10K SNP multiplex. Whole genome kinship cannot necessarily, and did not in this example, distinguish unrelated and related pairs since the overall number of “matching SNPs” is similar in both scenarios shown. Since windowed kinship uses locus proximity and searches for segments of shared kinship, it better distinguishes more distant relationships from 10K SNP data.

**Fig. S10:**
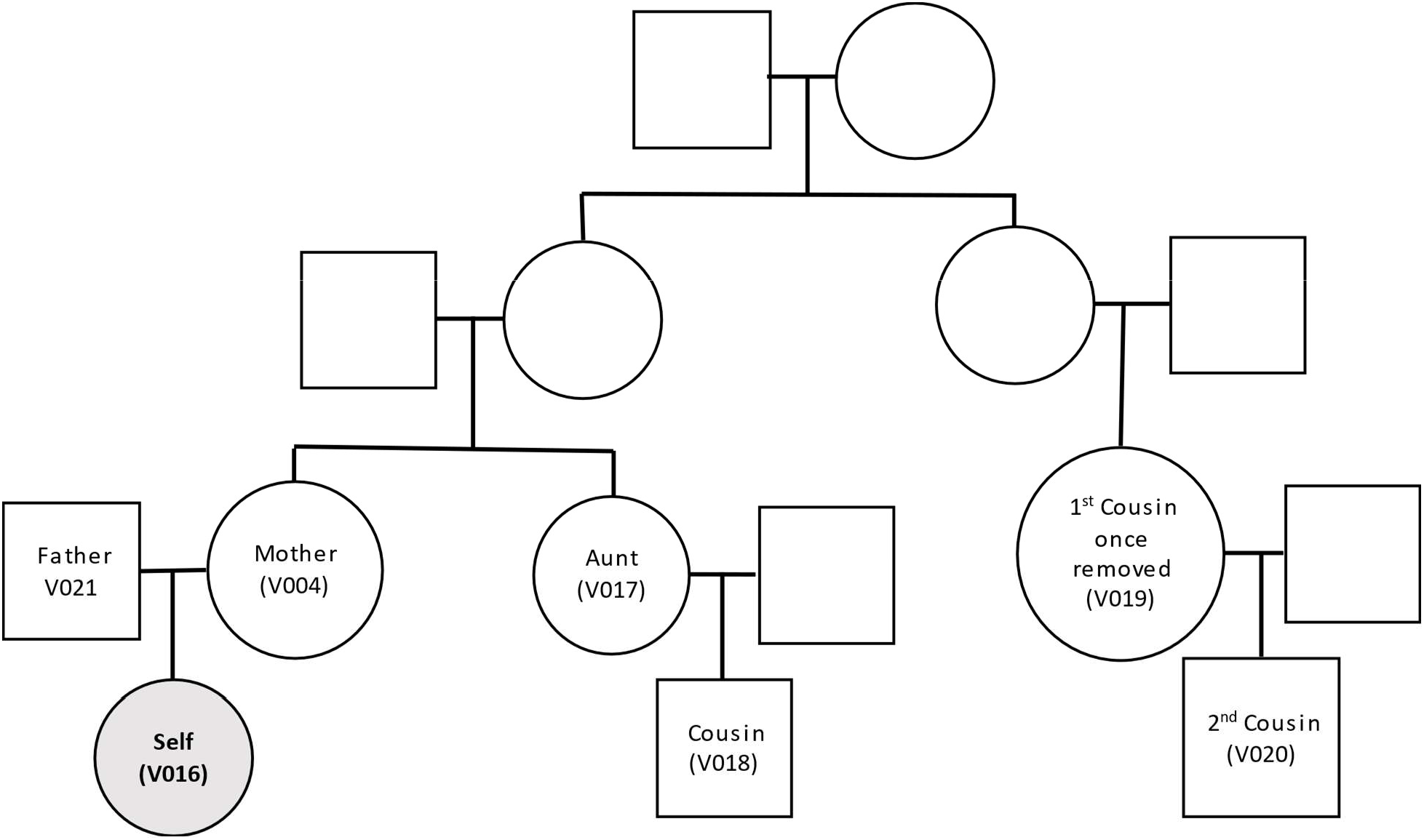
Known family pedigree used for mock casework study. All samples in labeled nodes were typed at the 10K SNP set using the ForenSeq Kintelligence Kit. Sample V016 was partially degraded, 250 pg template was used and read counts were reduced by increasing sample numbers per MiSeq FGx run to simulate a challenging case-type sample for FGG database query using the windowed kinship algorithm.

**Fig. S11:**
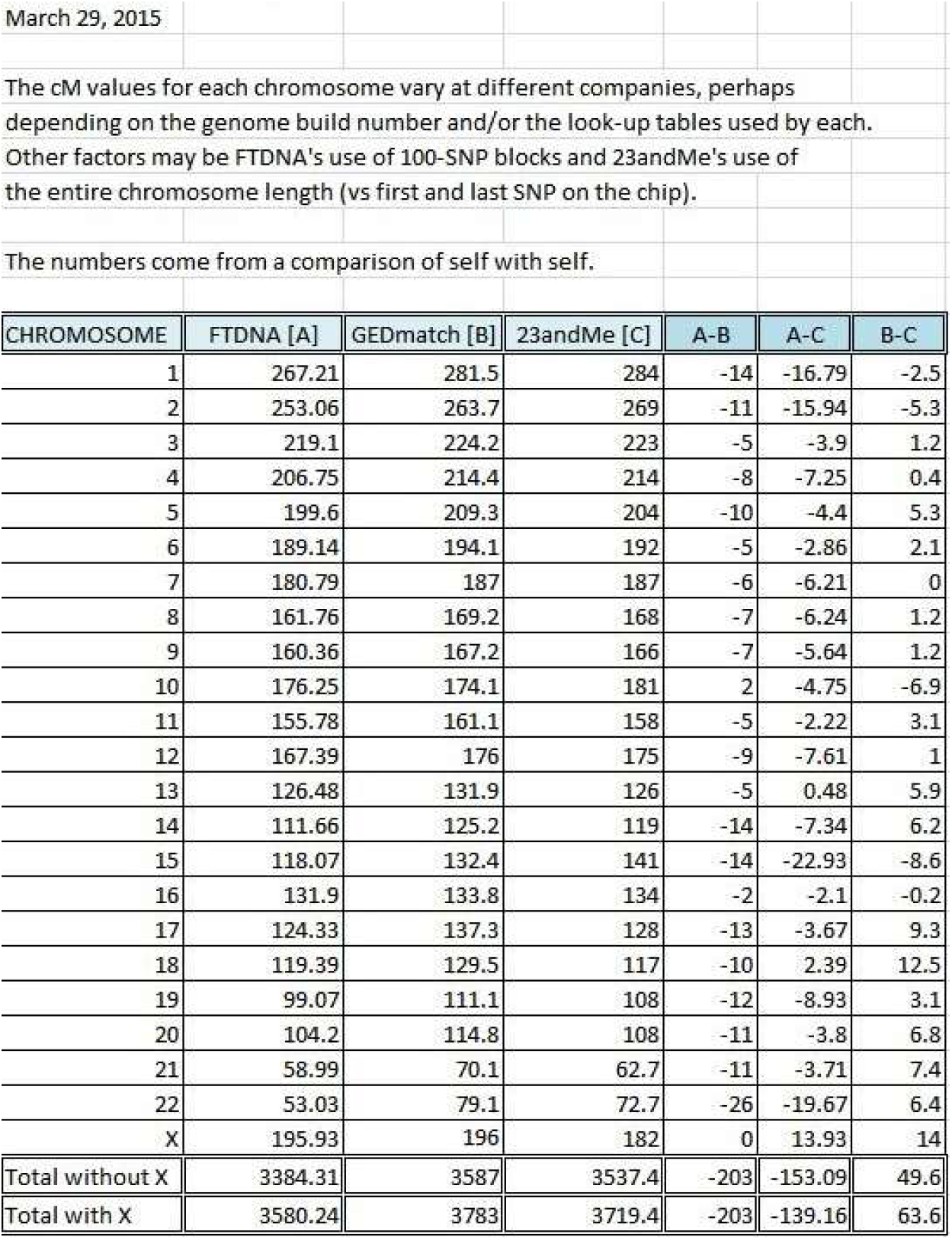
Comparison of total cM values per chromosome for FamilyTreeDNA, GEDmatch and 23andMe for one DNA sample. Sourced from ISOGG wiki.^7^

## Footnotes

1 Research purposes, in accordance with Terms of Service

2 https://www.gedmatch.com/

3 https://www.familytreedna.com/why-ftdna

4 https://thegeneticgenealogist.com/

5 https://classic.gedmatch.com/Documents/Qdocs.pdf

6 https://dnapainter.com/tools/sharedcmv4

7 https://isogg.org/wiki/CentiMorgan

